# Transcription decouples estrogen-dependent changes in enhancer-promoter contact frequencies and spatial proximity

**DOI:** 10.1101/2023.03.29.534720

**Authors:** Luciana I. Gómez Acuña, Ilya Flyamer, Shelagh Boyle, Elias Friman, Wendy A. Bickmore

## Abstract

How enhancers regulate their target genes in the context of 3D chromatin organization is extensively studied and models which do not require direct enhancer-promoter contact have recently emerged. Here, we use the activation of estrogen receptor-dependent enhancers in a breast cancer cell line to study enhancer-promoter communication. This allows high temporal resolution tracking of molecular events from hormone stimulation to efficient gene activation. We examine how both enhancer-promoter spatial proximity assayed by DNA fluorescence in situ hybridization, and contact frequencies resulting from chromatin in situ fragmentation and proximity ligation by Capture-C, change dynamically during enhancer-driven gene activation. These orthogonal methods produce seemingly paradoxical results: upon enhancer activation enhancer-promoter contact frequencies increase while spatial proximity decreases. We explore this apparent discrepancy using different estrogen receptor ligands and transcription inhibitors. Our data demonstrate that enhancer-promoter contact frequencies are transcription independent but are influenced by enhancer-bound protein complexes whereas altered enhancer-promoter proximity depends on transcription. Our results emphasize that the relationship between contact frequencies and physical distance in the nucleus, especially over short genomic distances, is not always a simple one.

## Introduction

How enhancers exert their action over long genomic distances is not understood. Contemporary models range from those that involve direct enhancer-promoter contact in 3D space, models that do not require enhancer-promoter contact with each other but do require engagement with transcriptional hubs, through to models based on diffusion of activating signals in a confined volume of the nucleus (Lim and Levine, 2021; Karr et al., 2022).

Two major methodologies – biochemical and imaging - are most often used to study the proximity of enhancers in relation to the gene promoters they regulate. Chromosome conformation capture (3C)-derived techniques rely on enzymatic fragmentation of cross-linked and detergent-treated chromatin followed by proximity ligation. When followed by next-generation sequencing, techniques such as Hi-C and micro-C can provide pair-wise contact frequencies genome-wide. Hundreds of thousands, to millions, of cells are often required to obtain high-resolution interaction maps, so the resulting data represent average contact frequencies of the cell population.

In imaging-based methods genomic loci are visualized either in live or fixed cells. Techniques, such as DNA fluorescence in situ hybridization (DNA-FISH) generally allow for analysis of only a handful of loci at a time in a few hundred cells but distances between loci in the nucleus can be determined at high spatial resolution at the single cell/allele level.

Generally, 3C- and imaging-based approaches give concordant views of 3D genome organization (Finn et al., 2019; Mateo et al., 2019; Boyle et al., 2020). However, there are examples where imaging data do not match the expectations from proximity ligation frequencies (Williamson et al., 2014; Finn et al., 2019), suggesting that the mechanistic basis for both data sets warrants further examination (Fudenberg and Imakaev, 2017).

A particularly fascinating area of dynamic 3D genome organisation is enhancer-promoter interaction. Hi-C and Micro-C data both show evidence for enriched interactions between enhancers and their target promoters in vertebrate genomes (Javierre et al., 2016; Hsieh et al., 2020). These data are consistent with a looping model of enhancer-promoter direct contact (Karr et al., 2022). However, a number of observations from imaging techniques are not consistent with a simple enhancer-promoter contact model where enhancers are juxtaposed to, or contact, their cognate target genes in a somewhat stable manner. Both live-cell imaging and DNA-FISH in fixed cells provide no evidence that very close proximity (<200nm) between enhancers and promoters is temporally correlated to enhancer-driven gene transcription in mouse embryonic stem cells (mESCs) (Alexander et al., 2019; Benabdallah et al., 2019; Platania et al., 2023). Indeed, DNA FISH indicates reduced enhancer-promoter proximity upon enhancer activation at the *Shh* locus (Benabdallah et al., 2019; Kane et al., 2022).

Enhancers in the complex regulatory landscapes of developmental genes may have different mechanisms of action from those that function in other biological contexts, such as in the acute responses to physiological cues. Estrogen-receptor alpha (ERα)-dependent enhancers in the human breast cancer cell line MCF-7 allow tracking of events after hormone treatment at high temporal resolution in a well-studied model system in which enhancers and their target genes, as well as many of the molecular mechanisms operating during enhancer activation, are well defined (Zwart et al., 2011). Chromosome conformation capture assays including 3C, Hi-C and ERα-selected CHIA-PET have been used to examine contact frequencies in MCF-7 cells before and after the addition of the ERα ligand 17β-estradiol (E2) (Fullwood et al., 2009; Hah et al., 2013; Rodriguez et al., 2019). By contrast, there has been little visual examination of the spatial proximity of specific ERα-enhancers and their target genes in the nucleus.

Here we use Capture-C and DNA-FISH in MCF-7 cells to examine the high-resolution 3D organisation of two genes whose transcription is induced rapidly and efficiently from ERα-bound enhancers upon E2 addition. We demonstrate that the two orthogonal methods produce seemingly paradoxical views of 3D enhancer-promoter organisation in response to induction and we explore the basis for these apparent discrepancies using different ligands of ERα and inhibitors of transcription. Our data demonstrate the need to carefully explore the role of enhancer-bound complexes, RNA polymerase II and transcription as determinants of 3C contact frequencies and the enhancer-promoter spatial distances determined by DNA-FISH.

## Results

### Rapid enhancer-dependent gene activation in response to estradiol

ERα is a member of the nuclear receptor transcription factor family; upon binding to the ligand E2 it binds chromatin at specific pre-marked regulatory regions (including enhancers) within minutes (Holding et al., 2018) inducing rapid transcription of target genes (Hah et al., 2011). Taking advantage of the extensive data available on transcription factor and co-activator chromatin binding in MCF-7 cells stimulated with E2, we identified two enhancer-gene pairs to study enhancer-promoter communication. *GREB1* and *NRIP1* are well studied E2 responsive genes (Hah et al., 2011). E2 responsive enhancers were identified using published ERα and p300 binding data from untreated and E2 treated cells (Hah et al., 2013; Li et al., 2013), and were paired to the target genes by proximity. A putative *GREB1* enhancer is located 40 kb upstream of the gene promoter and the *NRIP1* enhancer is located 100 kb upstream of the promoter (Fig. 1A).

**Figure 1.**
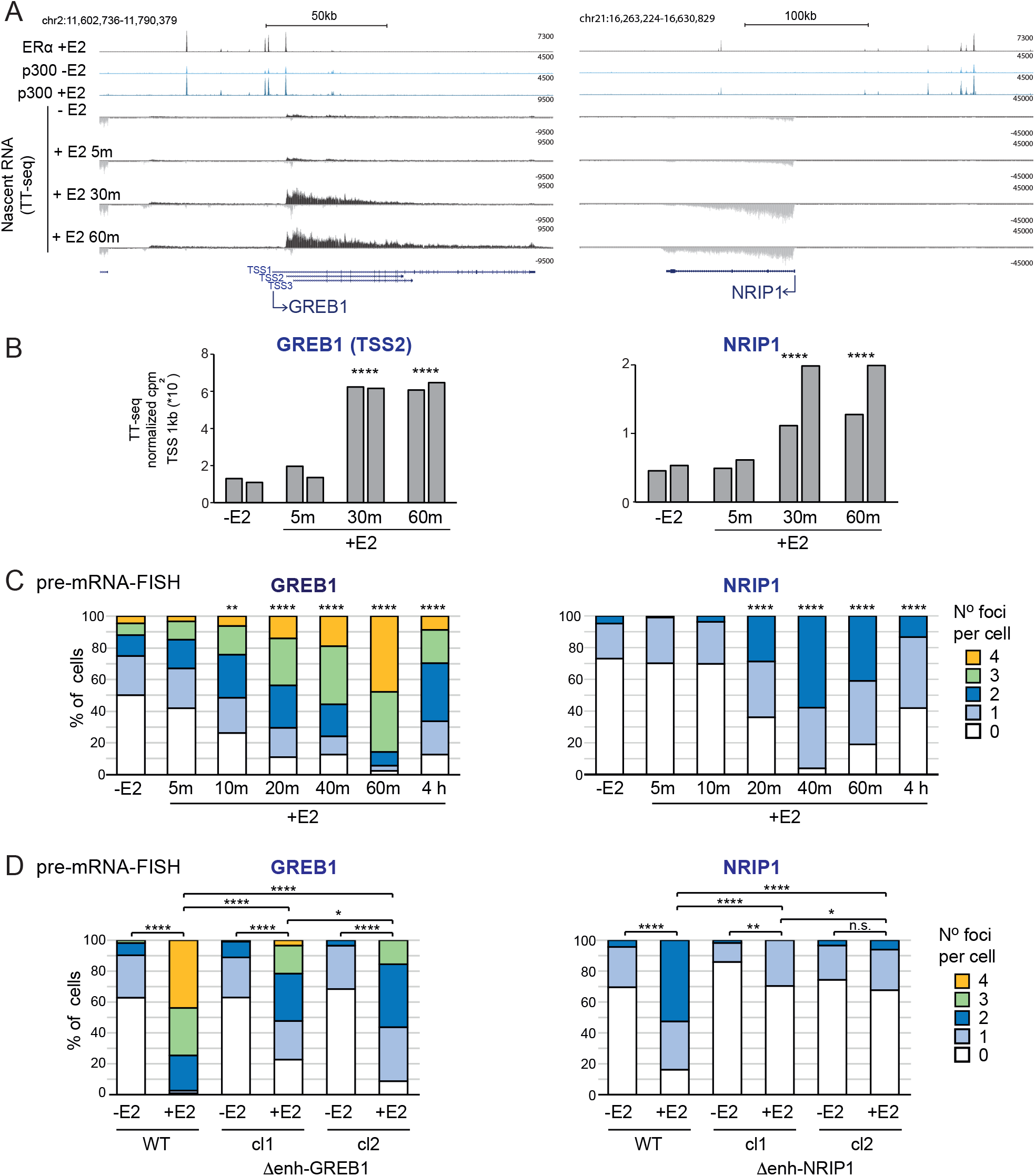
Rapid induction of ER-responsive genes. A) Genome browser snapshots of the *GREB1* (left) and *NRIP1* (right) loci showing published ChIP-seq tracks of ERα after E2 addition, p300 with and without E2 (Zwart et al, 2011) and TT-seq tracks without E2 and 5, 30 and 60 min after E2 addition in MCF-7 cells. Yellow bars indicate putative ERα enhancers. Genome coordinates: hg19 assembly of the human genome. B) Quantification of TT-seq reads over 1kb regions extending downstream of *GREB1* and *NRIP1* TSSs. Normalized counts per million reads (cpm) of two replicates are shown. C and D) pre-mRNA FISH for *GREB1* and *NRIP1* nascent transcripts without and with E2 in (C) MCF-7 cells for the indicated timepoints or (D) in cells where the respective putative ERα enhancers have been deleted. Results for two independent homozygous clones are shown. The percentage of cells with 0, 1, 2, 3, 4 foci is shown. Two-sided Fisher exact test. *p<0.05, **p<0.01, ***p<0.001, ****p<0.0001. Biological replicates for the data in panels C and D are in Supplementary Fig. S1. Statistical data for Figure 1 are in Supplementary Table S1.

Using nascent RNA-seq (TT-seq) (Schwalb et al., 2016), across a time course (5, 30 and 60 min) after E2 addition to MCF-7 cells which had been extensively starved of hormone, we detect abundant transcription from these genes 30 mins after E2 addition, but not at the earlier 5-minute time point (Fig. 1A, B; Supplementary Table S1). As previously reported using GRO-seq (Hah et al., 2013), TT-seq also reveals enhancer RNAs (eRNAs) transcribed at these ERα-bound sites and induced with similar kinetics to the gene mRNAs (Supplemental Fig. S1A, B; Supplementary Table S1). There seems to be a unidirectional long transcript running through the intergenic space upstream of *GREB1*, for *NRIP1* short eRNAs are detected at the sites of ERα-binding.

We confirmed these transcription dynamics at the level of individual alleles by nascent RNA-FISH using probes targeted to the first intron of each gene. There are 4 copies of *GREB1* in our MCF-7 cells - two copies on cytologically normal chromosomes 2, one on a chromosome with additional material translocated onto 2q and the fourth on a small portion of chromosome 2p translocated onto another chromosome (Kocanova et al., 2010). Pre-mRNA FISH detects significant upregulation of *GREB1* 10 min after hormone addition, with the frequency of active alleles increasing up to an hour after E2 addition when 79% of alleles are found to be active (Fig. 1C; Supplemental Fig.1C). *NRIP1* is on chromosome 21 of which there are 2 copies in MCF-7 cells. Significantly increased *NRIP1* transcription activation is observed after 20 min of stimulation, with levels reaching a maximum at 40 mins when 73% of alleles appear to be active (Fig. 1C; Supplemental Fig.1C; Supplementary Table S1). For both genes, the number of active alleles decreases by 4h after stimulation, but not down to the level of untreated cells.

The putative *GREB1* and *NRIP1* ERα enhancers have other features, such as p300 and FOXA1 occupancy (Hurtado et al., 2010; Hah et al., 2013), expected of a functional enhancer, but direct genetic evidence linking these sites to the activity of target genes has been lacking. To address this, we used CRISPR-Cas9 with guides targeting the borders of the ERα peaks to delete these putative enhancers. Two clones bearing homozygous deletions for each of the two enhancer regions were recovered (Supplemental Fig.1D). GREB1 enhancer deletion led to a significant decrease in the efficiency of *GREB1* activation in E2-treated cells detected by nascent RNA-FISH (Fig. 1D; Supplemental Fig.1E). Residual *GREB1* expression may result from multiple regulatory regions coordinating *GREB1* expression in E2 treated cells (Saravanan et al., 2020). Deletion of the NRIP1 enhancer led to almost total loss of *NRIP1* induction in response to E2 (Fig. 1D; Supplemental Fig.1E; Supplementary Table S1). In contrast, *NRIP1* was still highly E2 inducible in cells deleted for the GREB1 enhancer and *GREB1* E2-dependent induction was unaffected by deletion of the NRIP1 enhancer (Supplemental Fig.1E; Supplementary Table S1). These data confirm the role of the upstream ERα binding sites as *bona fide* enhancers driving E2-responsive transcription specifically at at *GREB1* and *NRIP1*.

### Increased enhancer-promoter contact frequency, but decreased spatial proximity, in response to estrogen

Whether enhancers drive expression of their target promoters through a direct physical interaction, or through contact-independent mechanisms is often assayed from the interaction frequencies derived from chromosome conformation capture assays (Leung et al., 2022). To investigate E2-dependent enhancer-promoter communication, we performed high resolution Capture-C (Golov et al., 2019), using tiled BACs covering a 500 kb region at *GREB1* and a 1 Mb region at *NRIP1,* in MCF-7 cells treated with vehicle, or with E2, at 5, 30 and 60 minutes. Relative enhancer-promoter contact frequencies increase upon E2 stimulation at both loci, with enhanced focal contacts detectable between the enhancer and promoter regions (Fig. 2A and Supplemental Fig. S2A, arrowheads). Using the enhancer regions as viewpoints in virtual 4C plots, increased contact frequency with the target promoter is detected as early as 5 min after E2 stimulation (Fig. 2B and Supplemental Fig 2B, arrowheads) and these enhancer-promoter contact frequencies are the highest in the whole captured regions (Supplemental Fig. S2C). Intriguingly, and most noticeable in the case of *GREB1*, transcription-related changes along the length of the gene body are also observed: while very short-range relative contact frequencies increase along the gene when it is transcribed, longer-range contact frequencies decrease (Fig. 2A and Supplemental Fig. S2A).

**Figure 2.**
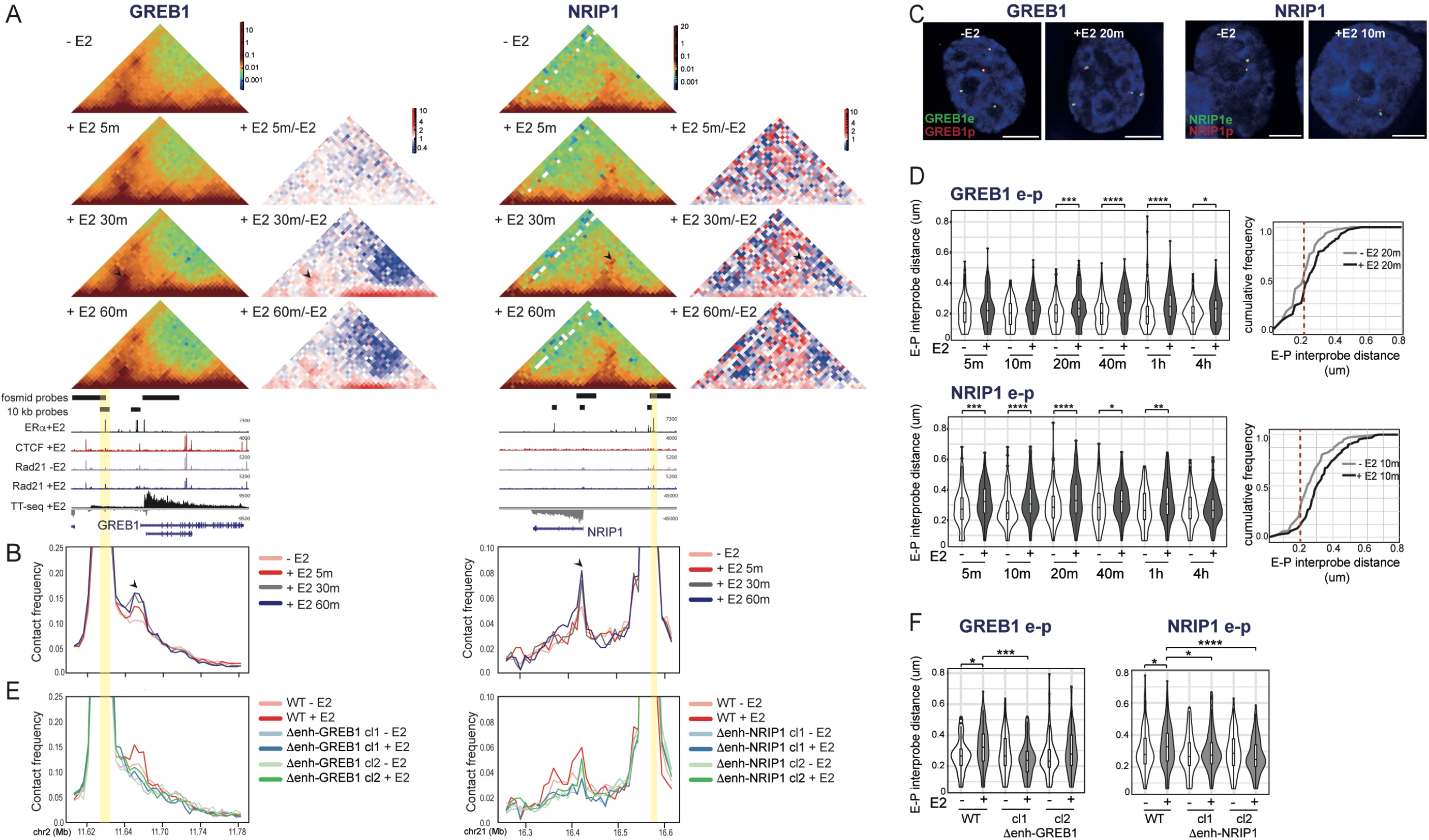
E2 induces increased Capture-C contact frequencies but decreased spatial proximity between enhancers and promoters. A) Capture-C heatmaps at *GREB1* (left; 5kb resolution) and *NRIP1* (right; 10kb resolution) loci from MCF-7 cells without (-E2) and with E2 (+E2) for the indicated time points. ERα +E2 (Zwart et al, 2011), CTCF +E2, Rad21 –E2, Rad21 +E2 ChIP-seq tracks (Schmidt et al, 2010), and TT-seq +E2 30 min are shown below. Enhancer regions are highlighted with yellow bars. In red-green-blue heatmaps (left), each pixel represents the normalized contact frequency between a pair of loci. In red-white-blue heatmaps (right) each pixel represents the ratio between E2-treated at the indicated time points and untreated samples. Blue indicates loss of contact frequency in treated vs untreated samples and red pixels indicate a gain. Arrowheads indicate pixels corresponding to enhancer-promoter pairs. Data from an independent biological replicate are in Supplementary Fig. S2A. B) Virtual 4C plots derived from the Capture-C normalized contact frequencies in (A) using the enhancer regions as viewpoints. Genome coordinates: hg19 human genome. Arrowheads indicate the promoters of *GREB1* and *NRIP1*. C) Representative images of nuclei (DAPI, blue) from untreated (-E2) or E2 treated MCF-7 cells showing DNA-FISH signal from fosmid probes targeted to the enhancer (e, green) and promoter (p, red) regions of the *GREB1* (left) or *NRIP1* (right) loci. The position of the probes is shown in (A). More detailed probe position is shown in Supplemental Fig. S2C. Scale bars, 5µm. D) Left: Violin plots showing the distribution of DNA-FISH inter-probe distances (μm) between e-p fosmid probe pairs at *GREB1* (top) or *NRIP1* (bottom) in untreated and E2 treated MCF-7 cells for the indicated time points. Boxes indicate median and interquartile distances. The statistical difference in data distribution +/-E2 at each time point was assessed by a two-sided Mann-Whitney test. *p<0.05, **p<0.01, ***p<0.001, ****p<0.0001. Holm-Bonferroni correction for multiple testing. Data from an independent biological replicate are in Supplemental Fig. S2D. Right: Cumulative frequency plots of the e-p inter-probe distances for the indicated time points and loci. Red dashed line indicates 200nm. E) As in (B) but for WT and enhancer deletion clones untreated or treated with E2 for 30 min. Heatmaps and data from an independent biological replicate are in Supplemental Fig. S4. F) Violin plots showing the distribution of DNA-FISH inter-probe distances between e-p fosmid probe pairs in WT and enhancer deletion MCF-7 clones untreated and E2 treated for 30 min. Data from an independent biological replicate are in Supplemental Fig. S4. Statistical data for Figure 2 are in Supplementary Table S2.

Imaging-based methods such as DNA FISH and live cell imaging have questioned whether enhancer-promoter juxtaposition is required for transcription (Benabdallah et al., 2019; Alexander et al., 2019; Kane et al., 2022), including for enhancer activation by nuclear hormone-dependent enhancers (Jubb et al., 2017). To explore this in the context of rapid gene induction by ERα-dependent enhancers, we used DNA-FISH with fosmid-derived probes spanning the promoter, enhancer and control regions at the *GREB1* and *NRIP1* loci (Supplemental Fig. S2C). We measured inter-probe distances at different time points after E2 stimulation. At both loci, enhancer-promoter inter-probe distances increase upon E2 treatment. In the case of *GREB1* this is statistically significant 20 min after stimulation, but the trend is already observable at 10 min (Fig. 2C,D; Supplemental Fig. S2D; Supplementary Table S2). At *NRIP1*, significantly increased enhancer-promoter spatial separation is observed as soon as 5 min after hormone stimulation, is sustained until 1 hour after E2 addition, but is no longer observed at 4h. Increased inter-probe distances are not seen between enhancers and control probe pairs (Supplemental Fig. S3A; Supplementary Table S2).

It is usually thought that regions showing high contact frequencies in 3C techniques fall within a 200 nm radius of each other and that, rather than looking at overall distance distributions and median values in imaging data, the proportion of loci at short distances is a more appropriate comparison (Fudenberg and Imakaev, 2017; McCord et al., 2020). However, despite the increased enhancer-promoter contact frequencies we detect by Capture-C after E2 addition, cumulative frequency analysis of inter-probe distances at both the *GREB1* and *NRIP1* regions show that the proportion of alleles with enhancer-promoter inter-probe distances < 200 nm decreases in E2 treated cells (Fig. 2D, right panels). No difference is observed in enhancer-control inter-probe cumulative distributions upon E2 addition (Supplemental Fig. S3A).

To ensure that any very focal enhancer-promoter co-localisation was not being obscured by our use of comparatively large (40kb) fosmid-derived probes, we repeated DNA FISH with probes detecting 10 kb regions centred on the sites with E2-triggered Capture-C higher contact frequencies. We also included a probe targeted to an intragenic (i) ERα ChIP peak *NRIP1* locus (Supplemental Fig. S2C). Twenty minutes after E2 addition, at both loci we still observe that enhancer-promoter distances increase, and the proportion of alleles at <200nm decreases, upon E2 addition (Supplemental Fig. S3B; Supplementary Table S2). Of note however, the absolute distances measured with the 10kb probes are smaller than those measured using the larger probe size. This may be due to to use of a confocal super-resolution imaging platform to image the signals from the smaller probes, providing improved resolution, especially in the *z* dimension, compared to the faster epifluorescence imaging platform used for all other experiments.

### E2-induced changes in enhancer-promoter contact frequency, and spatial proximity, are enhancer-dependent

To ascertain whether E2-induced changes in chromosome conformation and spatial organisation are dependent on specific ERα enhancer activation, we performed Capture-C and DNA-FISH on the cell clones deleted for either the *GREB1* or the *NRIP1* enhancers that we show are required for correct E2-dependent gene activation (Figs. 1D and Supplemental Fig. S1). The size of the engineered deletions - around 600 bp - is smaller than both the bin size used in our Capture-C analysis and the region covered by the DNA-FISH probes, making it possible to look at contact frequencies and spatial distances involving these deletion-harbouring regions.

At both *GREB1* and *NRIP1* loci, Capture-C shows that E2-triggered increased enhancer-promoter contact frequencies are lost upon enhancer deletion (Fig. 2E and Supplemental Fig. S4A,B arrowheads). Even though E2 induced *GREB1* transcription is not totally impaired by enhancer deletion, the impact of the decreased transcriptional activity on E2 dependent 3D chromatin structure within the *GREB1* transcribed unit is also seen in the Capture-C heatmaps (arrowed in Supplemental Fig. S4A). DNA-FISH shows that the E2 triggered increase in enhancer-promoter inter-probe distances is also lost in the enhancer-deleted cells at both studied loci (Fig. 2F; Supplemental Fig. S4C; Supplemental Table S2). The deletions had no impact on spatial distances measured between the enhancers and control probes (Supplemental Fig. S4D; Supplemental Table S2). We conclude that both E2-dependent contact frequencies, and enhancer-promoter spatial separation, are affected by the enhancer deletions but in opposite directions – a loss of enhancer-promoter contact in chromosome conformation capture assays and decreased enhancer-promoter spatial separation assayed by DNA-FISH.

### Enhancer-promoter contacts and spatial proximity are ligand dependent

Enhancer bound ERα recruits co-activators, including SRC1,2 and 3 and the histone acetyltransferases CBP and p300, as well as the transcriptional machinery (Zwart et al., 2011; Legare and Basik, 2016). Therefore, our data implicating liganded (E2) ERα binding in inducing enhanced enhancer-promoter “contact” and yet decreased spatial proximity, could result from the action of ERα itself, or the recruitment of additional transcriptional activators and co-activators.

Tamoxifen (4OH) is a selective estrogen receptor modulator widely used to treat ER-positive breast cancer. ERα liganded with 4OH has been shown to occupy broadly the same genomic sites as E2-ERα, including the *GREB1* and *NRIP1* enhancers (Fig. 3B) (Guan et al, 2019), where it recruits co-repressors such as NCOR, including histone deacetylases, repressing the hormonal response (Guan et al., 2019, Legare and Basik, 2016). Taking advantage of these differences in co-factor recruitment by E2 and 4OH we assayed the impact of 4OH treatment on chromatin conformation in MCF-7 cells. We verified ERα recruitment to chromatin after either E2 or 4OH treatment by immunofluorescence in detergent pre-extracted cells. Levels of chromatin-bound (detergent-resistant) ERα induced by 4OH are significantly higher than those observed with vehicle but are lower than with E2 (Supplemental Fig. S5A; Supplementary Table S3). Using RT-qPCR, we also verified that 4OH impedes transcriptional activation of E2-target genes. No induction of *GREB1* mRNA is detected by 4OH treatment (Fig. 3A; Supplementary Table S3). A low level of 4OH-induced *NRIP1* expression is detected, but significantly lower than that seen with E2.

**Figure 3.**
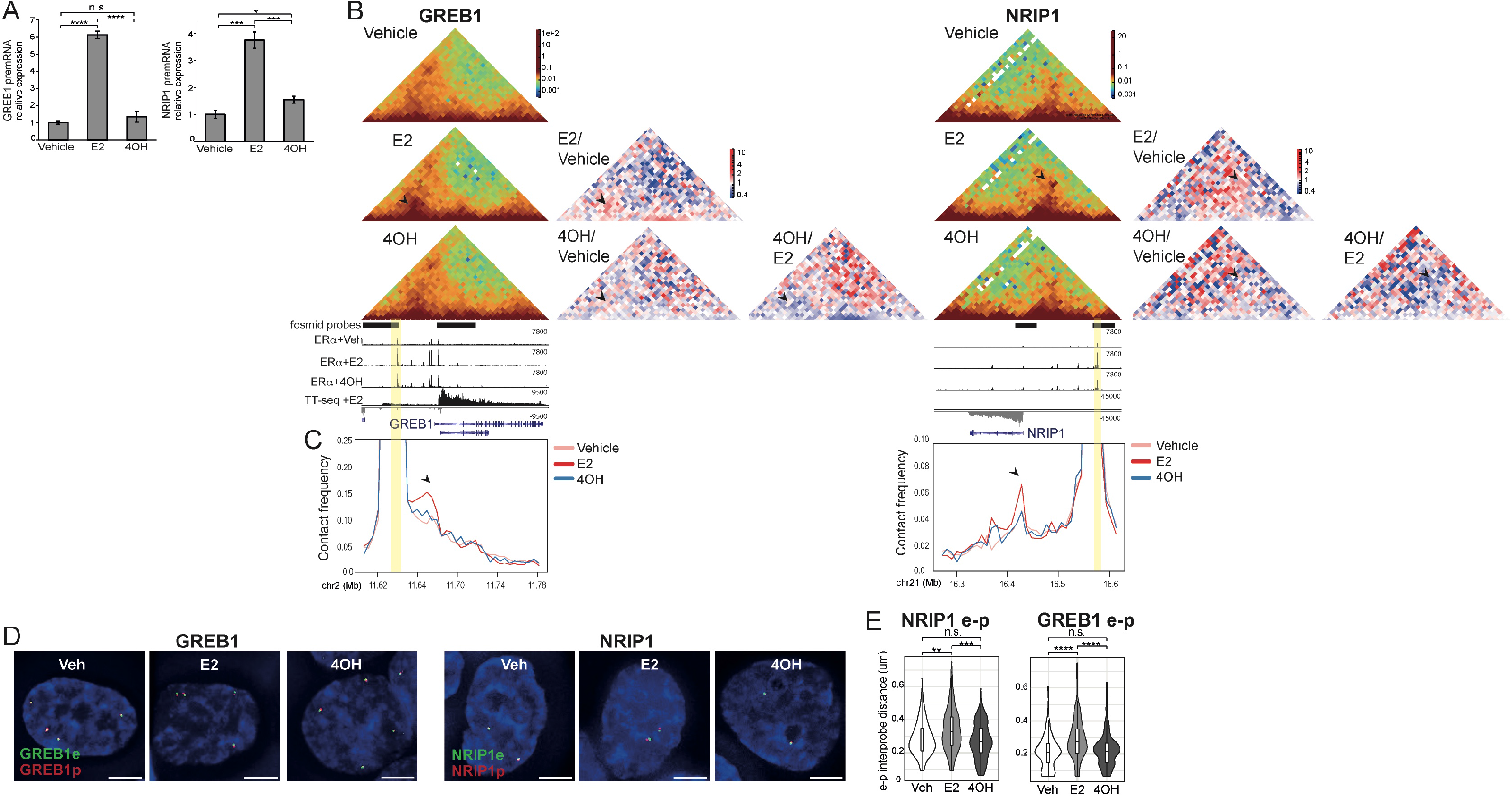
Enhancer-promoter interaction frequencies and spatial separation depend on the ERα ligand. A) RT-qPCR of *NRIP1* and *GREB1* pre-mRNAs in MCF-7 cells treated with vehicle, E2 or Tamoxifen (4OH) for 1h. T-test, Bonferroni correction for multiple testing. Mean +/- SD of 3 biological replicates. B) Capture-C heatmaps at *GREB1* (left; 5kb resolution) and *NRIP1* (right; 10kb resolution) loci from MCF-7 cells treated with vehicle, E2 or 4OH for 30 min. ERα +Vehicle, ERα +E2, ERα +4OH ChIP-seq data (Guan et al, 2019) and TT-seq +E2 30 min tracks are shown. Enhancer regions highlighted with yellow bars. In red-green-blue heatmaps, each pixel represents the normalized contact frequency between a pair of loci. In the red-white-blue heatmaps each pixel represents the ratio between 4OH-treated samples and vehicle- or E2-treated samples. Arrowheads indicate pixels corresponding to enhancer-promoter interaction frequencies. C) Virtual 4C plots derived from Capture-C normalized contact frequencies shown in (B) and using the enhancer regions as viewpoints. Arrowheads indicate the promoters. Genome coordinates: hg19. Data for a biological replicate are in Supplemental Fig. S5B,C. D) Representative images of nuclei (DAPI, blue) from cells treated with vehicle, E2 or 4OH and hybridised with fosmid probes targeted to the enhancer (e, green) and promoter (p, red) regions of *GREB1* or *NRIP1*. Scale bar, 5µm. E) Violin plots showing the distribution of DNA-FISH interprobe distances in cells treated with vehicle, E2 or 4OH using e-p fosmid probes at *GREB1* or *NRIP1* loci. Boxes indicate median and interquartile distances. Two-sided Mann-Whitney test, Holm-Bonferroni correction for multiple testing. n.s. p>0.05, **p<0.01, ***p<0.001, ****p<0.0001. Data from a biological replicate are shown in Supplemental Fig. S5D. Statistical data for Figure 3 are in Supplementary Table S3.

At the *GREB1* locus the enhancer-promoter contact frequencies induced by E2 (Fig. 2) are not seen with 4OH (arrowheads in Figs 3B, C and Supplemental Fig. S5B,C). Consistent with the absence of transcription in response to 4OH, the increased short-range contact frequencies along the length of the *GREB1* gene, that are induced in response to E2 (Fig. 2), are also absent in 4OH-treated cells (Fig. 3B; Supplemental Fig. S5B). Enhancer-promoter contact frequencies at *NRIP1* are also diminished in 4OH, relative to E2, treated cells (Fig. 3B,C; Supplemental Fig. S5B,C).

Unlike with E2, 4OH also does not induce any change in *GREB1* or *NRIP1* enhancer-promoter distances detected by DNA-FISH, compared with vehicle treated cells (Fig. 3D, E; Supplemental Fig. S5D; Supplementary Table S3). No changes in spatial distances between the *GREB1* or *NRIP1* enhancers and control probes are detected under any of the conditions (Supplemental Fig. S5E; Supplementary Table S3). We conclude that both enhancer-promoter contact frequencies and enhancer-promoter spatial proximity are not simply the consequence of ERα recruitment but reflect the nature of the co-factors and transcriptional machinery recruited thereafter.

### Spatial proximity, but not enhancer-promoter contacts, depend on transcription

To investigate the role of the transcriptional machinery in enhancer-promoter contacts and spatial proximity, we analysed the effects of RNA polymerase II (RPolII) inhibition on E2-induced changes in 3D chromosome conformation.

Flavopiridol inhibits CDK9/PTEF-b activity, RNA polymerase II Ser2 CTD phosphorylation and transcription elongation, whereas triptolide inhibits transcription initiation via TFIIH and leads to degradation of RPolII (Bensaude, 2011). We treated hormone starved MCF-7 cells with flavopiridol or triptolide for 5 min before adding E2 for 30 min (Fig. 4A). In this time frame we do not expect significant reduction of global RpolII levels (Jonkers et al., 2014, Chen et al., 2015). It was also reported that ERα, its co-activators, and the transcription machinery are still recruited to enhancers in the presence of flavopiridol, even though transcription of E2 target genes and eRNAs is inhibited (Hah et al., 2013). As expected, both inhibitors abolished the E2-dependent induction of *GREB1* and *NRIP1* mRNAs (Fig. 4A; Supplementary Table S4) but did not prevent the E2-dependant association of ERα with chromatin, though this is diminished in the case of triptolide (Supplemental Fig. S6A).

**Figure 4.**
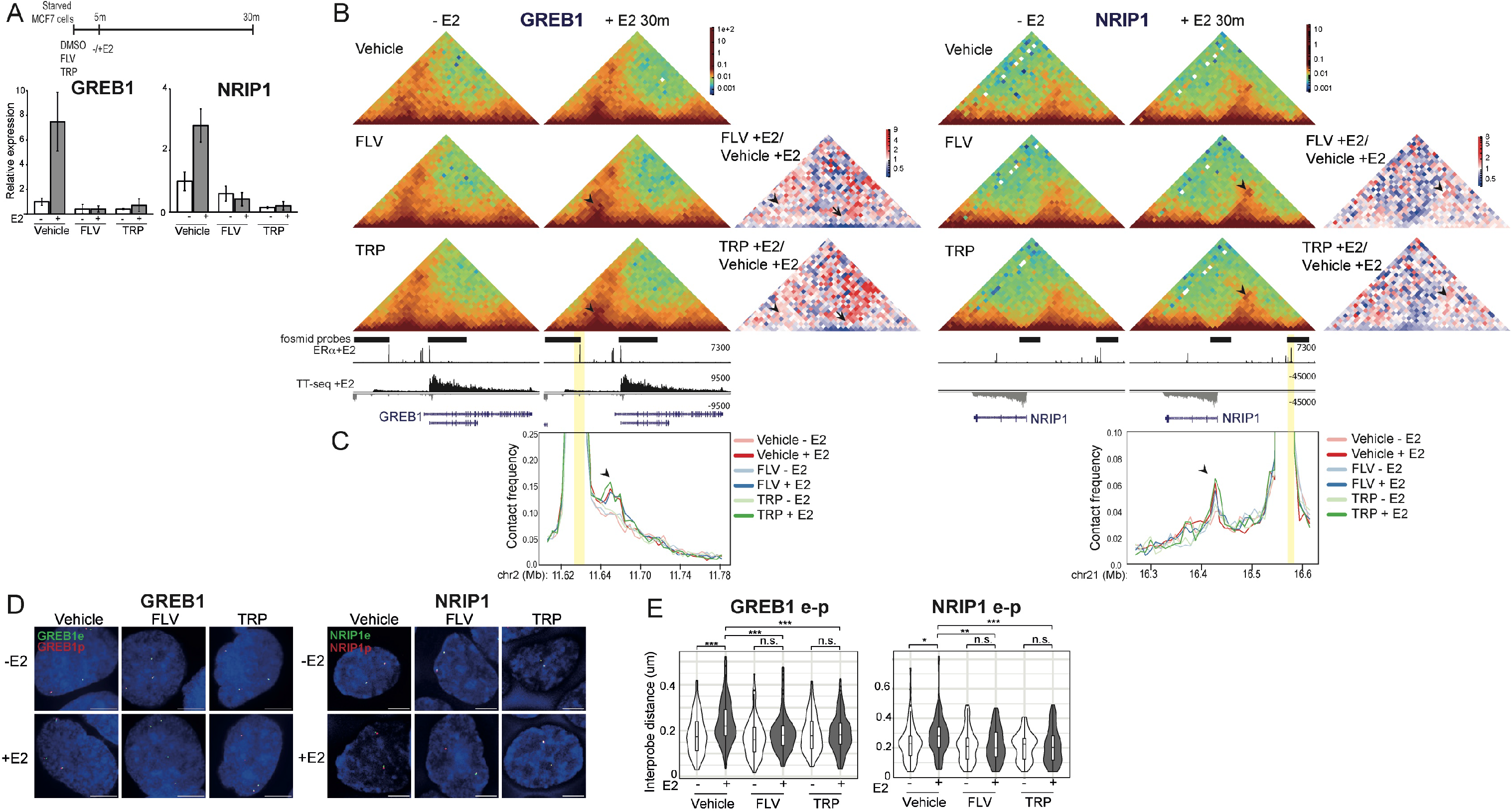
Inhibition of transcription impacts E2-induced enhancer-promoter spatial separation, but not Capture-C contact frequencies. A) Top; Schematic showing the treatment time-points, Below; RT-qPCR of *NRIP1* and *GREB1* pre-mRNAs in MCF-7 cells treated with vehicle, flavopiridol (FLV) or triptolide (TRP) for 5 min prior treatment without (-E2) and with E2 (+E2) for 1h. Means +/- SD of 3 replicates. B) Capture-C heatmaps at *GREB1* (5kb resolution) and *NRIP1* (10kb resolution) loci from MCF-7 cells treated with vehicle, FLV or TRP for 5 min prior treatment without (-E2) or with E2 (+E2) for 30 min. In red-white-blue heatmaps each pixel represents the ratio between FLV +E2 or TRP +E2, and vehicle +E2 treated samples. Arrowheads indicate pixels corresponding to enhancer-promoter interaction frequencies. Data from an independent biological replicate are in Supplemental Fig. S6B. C) Virtual 4C plots derived from the Capture-C normalized contact frequencies shown in (B) using the enhancer regions as viewpoints. Genome coordinates: hg19. Arrowheads indicate the promoters. D) Representative images of nuclei (DAPI, blue) of MCF-7 cells treated as in (A) showing the DNA-FISH signal from fosmid probes targeted to the enhancer (e, green) and promoter (p, red) regions of either GREB1 or NRIP1 loci. Scale bars, 5µm. E). Violin plots showing the distribution of DNA-FISH interprobe distances of cells treated as in (A), using e-p fosmid probes for either GREB1 or NRIP1 loci. Boxplots indicate median distances. Two-sided Mann-Whitney test, Holm-Bonferroni correction for multiple testing. Data from a biological replicate are in Supplemental Fig. S5D. n.s. p>0.05, *p<0.05, **p<0.01, ***p<0.001. Statistical data for Figure 4 are in Supplementary Table S4.

E2 also still induced increased Capture-C enhancer-promoter contact frequencies at both *GREB1* and *NRIP1* in the presence of flavopiridol and triptolide (Fig. 4B, C arrowhead and Supplemental Fig. S6B,C), but consistent with the inhibition of transcription, both inhibitors blocked the E2-induced gain of short-range contacts across the *GREB1* gene body.

In contrast, flavopiridol and triptolide abolished the E2-induced loss of enhancer-promoter spatial proximity at *GREB1* and *NRIP1*, as assayed by DNA FISH (Fig. 4D, E; Supplemental Fig. S6D; Supplementary Table S4). There was no effect of E2 on the distances between control probes under all conditions (Supplemental Fig. S6E; Supplementary Table S4). The use of these two RPolII inhibitors therefore allows us to separate the effects of E2 addition on 3D genome organisation as assayed by chromosome conformation capture or DNA FISH: while enhancer-promoter contact frequencies assayed by Capture-C are not affected by transcription inhibition, the enhancer-promoter spatial separation observed by DNA-FISH is transcription-dependent.

## Discussion

Chromosome conformation capture and 3D DNA-FISH are linchpins methodologies for 3D genome investigation. For the most part data from these orthogonal approaches are congruent, but in a few cases spatial distances measured by DNA-FISH and proximity ligation frequencies from chromosome conformation capture between the same loci are not so readily reconciled (Williamson et al., 2014; Finn et al., 2019).

One of the most intensively studied areas of 3D genome organisation is the relationship between enhancers and their target gene promoters. Whereas high-resolution chromosome conformation capture studies show enriched enhancer-promoter contact-frequencies (Javierre et al., 2016; Hsieh et al., 2020), there are several examples where live-cell imaging and 3D DNA-FISH in mammalian cells do not show the frequent enhancer-promoter proximity (<200nm) that might be expected from the results of proximity ligation-based assays (Alexander et al., 2019; Benabdallah et al., 2019; Platania et al.,2023). Contact-independent models are emerging that require general spatial enhancer-promoter proximity but not protracted enhancer-promoter direct contact for the activation of transcription (Kane et al., 2022; Karr et al., 2022; Rinzema et al., 2022).

Inducible systems are key to understanding how enhancer-promoter proximity is linked to gene activation. In mESCs, synthetic transcription factors have been used to study enhancer-driven gene activation unexpectedly revealing decreased enhancer-promoter spatial proximity upon enhancer activation (Benabdallah et al., 2019, Kane et al., 2022), inconsistent with an enhancer-promoter direct contact looping model (Karr et al., 2022). However, both the efficiency of transcription and the temporal resolution in these systems are low, such that events occurring immediately downstream of enhancer activation, and the genome organisation specific to transcribing alleles, may have been missed.

Here we address these limitations using a system of rapid and efficient enhancer-dependent gene activation in MCF-7 cells mediated by the binding of liganded ERα at the enhancers of two well studied target genes – *GREB1* and *NRIP1*. Upon hormone stimulation, ERα rapidly binds to the enhancers, which are already in an open chromatin state and bound by pioneer transcription factors (Holding et al., 2018, Glont et al., 2019). This ensures rapid and highly efficient transcriptional induction of target genes – nascent RNA FISH showed that the majority of alleles are in the process of being transcribed 20-60 mins after E2 addition (Fig. 1). Data on 3D genome organisation assayed in the E2-stimulated state is therefore chiefly attributable to actively transcribing alleles.

Using Capture-C we detected the increased enhancer-promoter contact frequencies upon hormone stimulation expected from enhancer-promoter looping models. However, DNA-FISH reveals a loss of enhancer-promoter spatial proximity at these same time points (Fig. 2), consistent with what we have seen at a developmental locus (Benabdallah et al., 2019, Kane et al., 2022), and at target loci of another nuclear hormone receptor – the glucocorticoid receptor (Jubb et al., 2017). We note that this contrasts with live cell imaging data in Drosophila embryos in which enhancer-driven transcription of a reporter gene corresponds to more sustained spatial proximity between the reporter and enhancer loci (Chen et al., 2018).

We used genetic and biochemical perturbations to dissect the molecular events driving these apparently opposing observations. Both the E2-stimulated increased enhancer-promoter Capture-C contact frequencies, and the decreased spatial separation detected by DNA-FISH, depend on the presence of the enhancers (Fig. 2). This contrasts with the lack of significant enhancer-promoter distance changes measured at the *Sox2* locus in mESCs when the SCR enhancer is mutated compared to wild-type cells (Platania et al., 2023).

Both the E2-stimulated increased enhancer-promoter Capture-C contact frequencies, and the decreased spatial separation, also depend on liganded ERα recruiting co-activator complexes and/or components of the transcription machinery, since they are lost in the presence of the ERα antagonist tamoxifen (Fig. 3). Finally, the use of transcription inhibitors allowed us to further disentangle the drivers of enhancer-promoter contact frequencies and spatial proximity. While E2-dependent enhancer-promoter contact frequencies persist when transcription initiation or elongation are blocked with triptolide or flavopiridol, the E2-dependent loss of enhancer-promoter proximity seen by DNA-FISH is abrogated (Fig. 4). In the time frame of the experiments performed here, ERα - along with co-activators and the transcription machinery, is known to be still recruited to chromatin by E2 in the presence of these inhibitors (Jonkers et al., 2014, Chen et al., 2015, Hah et al., 2013). It was reported that transcription inhibition has little impact on higher order 3D chromatin organization in mESCs (Hsieh et al., 2020) while enhancer-promoter contact frequencies were shown to be somewhat reduced (Hsieh et al., 2020, Barshad et al., 2023). It should be noted however, that this reduction is only noticeable at the genome wide level and that the foci of these interactions were largely unaffected (Hsieh et al., 2020, Barshad et al., 2023). Our observations also appear to differ from what has been seen at the Sox2-SCR locus in mESCs where triptolide has no affect on measured enhancer-promoter differences (Platania et al., 2023). However a notable difference is that whereas *Sox2* is being actively transcribed in mESCs, here we are looking at the acute transcriptional induction of genes from an inactive state.

We conclude that the gain of enhancer-promoter contact frequencies detected by Capture-C coincides with E2-liganded ERα recruitment to enhancers, whereas decreased enhancer-promoter proximity corresponds with E2-dependent transcription. Most noticeably in the case of *NRIP1*, increased enhancer-promoter separation occurs as soon as 5 min after E2 stimulation. This is before nascent transcription of the gene is detected by either TT-seq or RNA FISH (Fig. 1). However some, albeit not statistically significant, transcription of eRNAs is detected at this early timepoint (Supplemental Fig. S1).

We also detected an enrichment of short-range, and depletion of longer-range, Capture-C interactions along gene bodies linked to E2-induced transcription,. These depend on transcription, being lost in the presence of tamoxifen or transcription inhibitors. This is most noticeable in the case of *GREB1*, a long gene with approximately 30 exons, in comparison with *NRIP1’s* simple gene structure (up to 4 exons). These observations are compatible with a model in which genes stiffen and decompact as the transcription machinery elongates through, and nascent RNAs and RNA binding proteins coat, the gene bodies constraining their ability to make longer-range contacts (Leidescher et al., 2022).

At the cell population level, we observe increased Capture-C contact frequencies between regions that, at the same time, appear to become further apart, on average within the nucleus. Whilst imaging data is collected from almost every allele in the few hundred cells that are imaged per experiment, it is not know what proportion of alleles in the millions of cells that go into the Hi-C reaction contribute to the enhancer-promoter contact frequencies detected in Capture-C. We therefore cannot exclude that there is very transient physical contact between the enhancers and promoters that is too infrequent to detect by DNA-FISH, but that is captured by Capture-C. However, the fact that most of the alleles we visualise in this system are being actively transcribed means that such a “kiss and run” contact is not required for each burst of transcription.

Both 3C techniques and DNA-FISH rely on protein-protein and DNA-protein crosslinking with formaldehyde, which is widely used and yet incompletely understood (Hoffman et al., 2015). The recruitment of ERα, co-activators and the transcriptional machinery results in megadaltons of potentially additional cross-linkable protein complexes at ER-responsive enhancers following E2, but not 4OH, treatment. Crosslinking between somewhat distant genomic regions is also plausible via a crosslinked meshwork of macromolecules that bridge between two loci. This crosslinking radius is thought to depend on protein occupancy, the abundance of specific amino acids and crosslinking efficiency (Gavrilov et al., 2015; Giorgetti et al., 2016). Indeed, we note that the physical enhancer-promoter distances we measure upon E2 stimulation might fall within estimated 3C crosslinking and ligation radii (Giorgetti et al., 2016, Finn et al., 2019).

Formaldehyde crosslinking is also affected by protein dynamics. Protein cross-linking to DNA requires a long residence time (many seconds) (Schmiedeberg et al., 2009; Teves et al., 2016) and proteins can artifactually concentrate into large puncta, or can appear to leave foci, during fixation if the fixation rate is slower than protein diffusion rates (Irgen-Gioro et al. 2022). E2-dependent increases in enhancer-promoter contact frequencies detected by Capture-C could therefore be influenced by the residence time and exchange rates of the large multicomponent protein complexes recruited by ERα (Fig. 5). In contrast, the loss of enhancer-promoter spatial proximity detected by DNA-FISH appears to be linked to the production of RNAs (eRNA and mRNA) at the target loci, perhaps due to the effect of these RNAs on biomolecular condensate formation (Lee et al., 2021).

**Figure 5.**
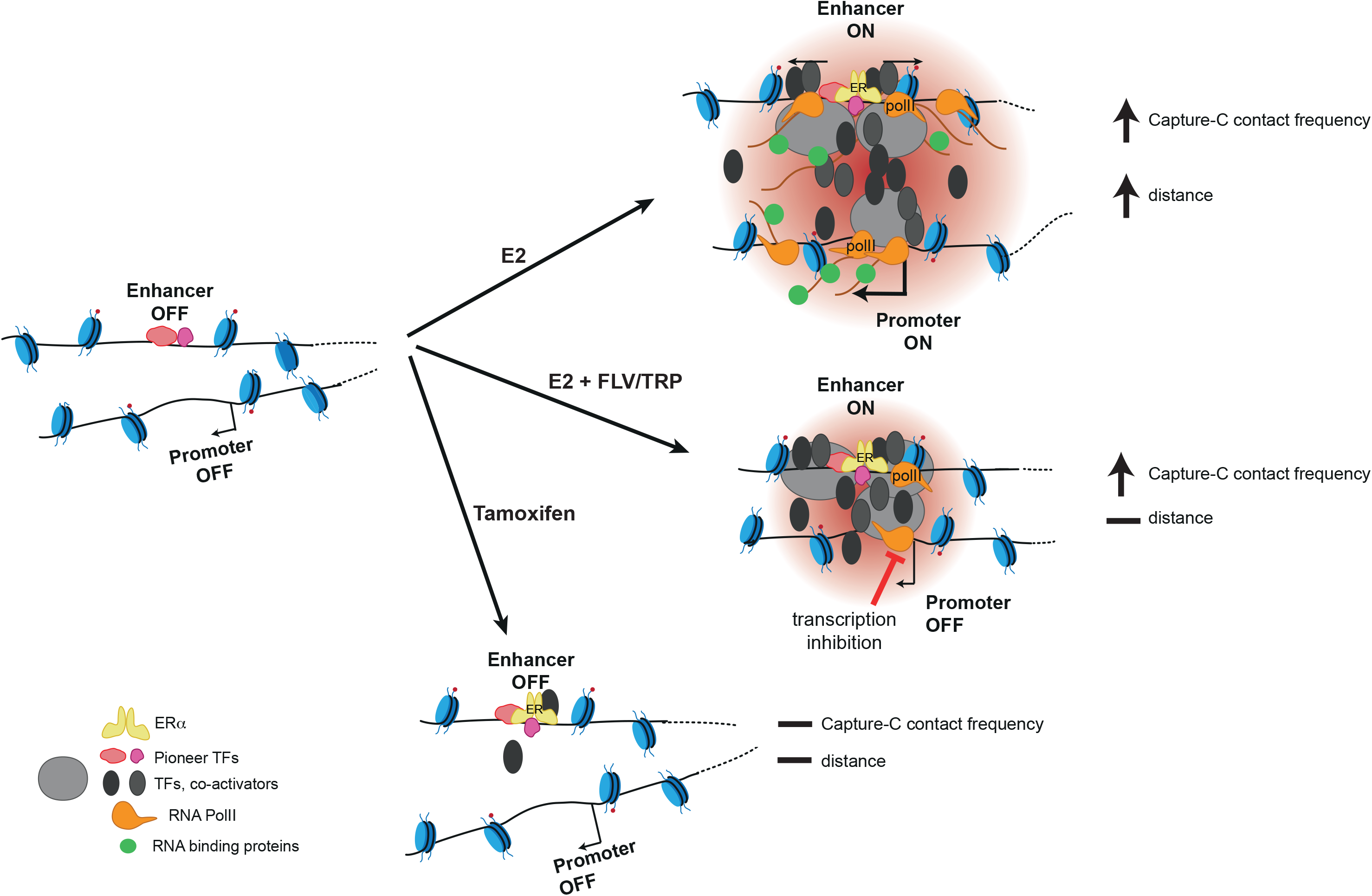
Model for enhancer-promoter communication leading to increased Capture-C contact frequencies but loss of spatial proximity during transcriptional activation. Upon estradiol (E2) stimulation, ERα binds to enhancers recruiting co-activators and the transcription machinery, and leading to the transcription of eRNAs and target mRNAs. This produces increased enhancer-promoter contact frequencies and yet increased spatial separation of enhancer and promoter (top right). When transcription is inhibited with flavopiridol (FLV) or triptolide (TRP) during E2 stimulation, ERα, co-activators and the transcription machinery are still recruited but without the transcription of eRNAs or mRNAs. This still produces increased enhancer-promoter contact frequencies without changes in spatial distance (middle right). With tamoxifen as ligand, ERα binds to the enhancers but recruits co-repressors, thus not inducing the activation of the target genes. In this context, neither enhancer-promoter contact frequencies nor spatial distances change in comparison with the unstimulated situation (bottom right). We speculate that the increase in enhancer-promoter contact frequencies observed upon hormone stimulation is driven by the recruitment of large multicomponent protein compexes while increased enhancer-promoter spatial separation seems to linked to the molecular events associated with transcription.

Our work reinforces that high resolution enhancer-promoter contact frequencies derived from 3C techniques cannot be simply interpreted as physical distances. Further careful consideration needs to be given to how dynamics and the functional, biochemical and biophysical environments of genomic loci in different biological systems and under different conditions may impact the data obtained from imaging and chromosome conformation assays.

## Methods

### Cell culture and treatments

MCF-7 cells were grown in Dulbecco’s Modified Eagle’s medium (DMEM, GIBCO021969-035), high glucose and supplemented with 2 mM L-glutamine, 100 units/ml penicillin, 6.5 μg/ml streptomycin and 10% foetal calf serum (FCS, Gibco 10270).

Before hormone stimulation, cells were seeded in growth media and 24hr later transferred into starving media comprising phenol-red free DMEM, penicillin/ streptomycin and 10% charcoal-stripped FCS for 96hr. Cells were transferred into phenol-red free DMEM supplemented with either 10 nM 17ß-estradiol (Sigma E2758) (+E2) or ethanol (-E2) for the indicated time points. 100 mM Tamoxifen (Sigma T3652) (4OH) was used. For transcription inhibition, media was supplemented with either DMSO (vehicle), 10 μM Flavopiridol (Cambridge Bioscience 2090-2) or 10 μM Triptolide (Sigma T3652). After 5 mins, 1/30 vol of media was incorporated with either ethanol (-E2) or 17ß-estradiol (+E2) to a final concentration of 10 nM for 30 mins.

### TT-seq

TT-sequencing was adapted from Boyle et al. (2020). 3x10^6^ MCF-7 cells were seeded in T75 flasks, starved, and then treated with E2 or vehicle for the indicated time points. For every time point, 4-thiouridine (4SU; Sigma T4509) was added to a final concentration of 500 µM for 5 min before cells were harvested with 2ml/flask of Trizol (Invitrogen 15596026). RNA was prepared as previously described (Boyle et al.,2020). RNA samples were sonicated with a Diagenode Bioruptor Plus for 30 s at a high setting. 10-100 ug of RNA were biotinylated, uncoupled biotin was removed and biotin-labelled RNA purified as previously descried (Boyle et al., 2020). Libraries were constructed using the NEBNext Ultra II directional RNA library preparation kit for Illumina according to the protocol for purified mRNA or ribosome-depleted RNA and with a 5-min RNA fragmentation step (NEB E7760). Library PCRs were supplemented with 2× SYBR dye (Sigma S9430) to monitor amplification by quantitative(q)PCR on a Roche LightCycler 480. To allow for sample multiplexing, PCRs were performed using index primers (NEBNext multiplex oligos for Illumina, set 1, E7335) and amplified to linear phase. Libraries were combined as an equimolar pool and sequenced on an Illumina NextSeq on a single high-output flow cell (paired-end 75-bp reads).

### TT-seq data analysis

For each demultiplexed sample, multiple raw Fastq files were merged (individually for reads 1 and 2). Adapter sequences were removed using Cutadapt v1.16 in paired end mode (options: -q 20 -a AGATCGGAAGAGCACACGTCTGAACTCCAGTCA -A AGATCGGAAGAGCGTCGTGTAGGGAAAGAGTGT -m 20). Adapter-trimmed Fastq files were aligned to the human genome (hg19) using STAR v2.7.8 for paired end sequence data and using Ensemble GRCCh37gene annotation to generate BAM files (options: --outFilterType BySJout -- outFilterMultimapNmax 20 --outSAMunmapped Within). Bam files were indexed using samtools (v1.3) index; bigwig files used for data visualization on UCSC genome browser, one file per strand, were generated using deeptools (v3.5.1) bamCoverage (options: --normalizeUsing RPKM -bs 1 - -filterRNAstrand forward or reverse). Replicates were merged in single bigwig file using deeptools bigwigCompare using the mean operation. Read counts over specific genomic regions were computed using featureCounts v2.0.1 (options: -p -Q 10 -s 2). To compute read counts over TSSs, regions 1kb downstream of TSSs of Ensemble GRCCh37gene annotation were used. To compute read counts over intergenic regions, pooled ATAC-seq peaks (Guan et al., 2019, see below for methods) at least 1kb away from annotated TSSs and extended 1.5 kb at each side, were used. Differential expression analysis was performed using edgeR on R.

### ChiP-seq data analysis

Data from Zwart et al. (2011) (ArrayExpress: E-MTAB-785), Schmidt et al (2010, ArrayExpress: E-TABM-828) and Guan et al (2019, NCBI GEO: GSE117943) were re-analyzed. Fastq files were quality assessed (FastQC v0.11.4), adapters trimmed (Cutadapt v1.16) and aligned to the human genome (hg19) using Bowtie2 v2.2.6 in either single- or paired-end modeto generate sam files (options for paired-end: --no-discordant --no-mixed --no-unal --very-sensitive -X 2000). Potential PCR duplicates were removed using samtools (v1.3) rmdup. Bigwig files used for data visualization on the UCSC genome browser were generated using deeptools (v3.5.1) bamCoverage (options: --normalizeUsing RPKM -bs 1). Replicates were merged in single bigwig file using deeptools bigwigCompare using the mean operation. Peaks were called using MACS2 (v2.1.1) callpeak (options: -g hs -B -q 0.01) for each replicate. Common peaks for each treatment were obtained using bedtools (v2.27.1) intersect.

### RNA-FISH

Custom Stellaris RNA FISH probes were designed against *GREB1* and *NRIP1* nascent mRNA first introns (pool of 48 unique 22-base polymer probes) using the Stellaris RNA FISH Probe Designer (www.biosearchtech.com/stellarisdesigner (version 4.2)). 4x10^5^ MCF-7 cells were seeded on Thermo Scientific SuperFrost Plus Adhesion slides (15438060), starved and treated as specified. Cells were fixed with 4% paraformaldehyde (pFa) in PBS for 10 min, rinsed with PBS 3 x for 2 min and permeabilized with 0.5% Triton X-100/ PBS for 10 min. Following 3 x 2 min rinses, slides were incubated in wash buffer (2× SSC, 10% deionized formamide) for 5 mins at r.t. Slides were hybridized with *GREB1* or *NRIP1* Stellaris FISH Probe sets labeled with Quasar 670 (Biosearch Technologies) following the manufacturer’s instructions (www.biosearchtech.com/stellarisprotocols). RNA FISH probes were hybridised to slides as previously described (Kane et al., 2022).

### 3D DNA-FISH

#### Probe preparation

For fosmid probes (Supplementary Table S6), 1 μg of fosmid DNA was labeled by nick translation to incorporate biotin-dUTP (Roche 11093070910) or digoxigenin-dUTP (Roche 11093088910) and prepared for hybridisation as previously described (Boyle et al., 2020). For 10kb PCR probes (Supplementary Table S6), fragments generated by PCR from BAC templates (Supplementary Table S7) were cloned into TOPO-TA plasmids (Invitrogen). 1 μg of plasmid DNA was labeled by nick translation to incorporate biotin-dUTP (Roche 11093070910), digoxigenin-dUTP (Roche 11093088910) or Green496-dUTP-labeled (Enzo Life Sciences) and used as above.

#### Cell fixing, denaturing and hybridization

Cells were seeded, starved, treated, fixed and permeabilized as for RNA-FISH. Following permeabilization and rinsing, slides were air dried and stored at -80°C till use. Slides were incubated in 100 μg/mL RNase A in 2× SSC for 1 hr at 37°C, washed briefly in 2× SSC, passed through an ethanol series (70, 90 and 100%, 2 min in each), and air-dried. Slides were incubated for 5 min in a dry incubator at 70°C, denatured in 70% formamide/2× SSC (pH 7.5) for 40 min at 80°C, cooled in 70% ethanol for 2 min on ice, and dehydrated by immersion in 90% ethanol for 2 min and 100% ethanol for 2 min prior to air drying. Probes in hybridization mix were added to the slides and incubated o/n at 37 °C. Following a series of washes in 2× SSC (45 °C) and 0.1× SSC (60 °C), slides were blocked in blocking buffer (4× SSC, 5% Marvel) for 5 mins. For 2-color detection (either with fosmid or 10 kb derived probes), the following antibody dilutions were made: anti-digoxigenin-Rhodamine FAB fragments (Roche, cat. no. 11207750910) 1:20; Texas Red anti-sheep 1:100 (Abcam ab6745) and Avidin-FITC1:500 (Vector Laboratories A-2011); biotinylated anti-avidin (Vector Laboratories, cat. no. BA-0300) 1:100, and Avidin-FITC 1:500. For 3-color detection the following antibody dilutions were made: anti-digoxigenin (Roche, cat. 11333089001) 1:10; AF647 anti-seep (Invitrogen, cat. A-21448) 1:10 and Texas Red avidin (Vector Laboratories, cat. no. A-2016) 1:500; biotinylated anti-avidin (Vector Laboratories) 1:100; Texas Red avidin (Vector Laboratories, cat. no. A-2016) 1:500. Slides were incubated with antibody in a humidified chamber at 37 °C for 30–60 mins in the following order, with 4× SSC/0.1% Tween 20 washes in between: fluorescein anti-digoxigenin, fluorescein anti-sheep, biotinylated anti-avidin, streptavidin-Cy5 for 3-color detection; Texas Red avidin, biotinylated anti-avidin, Texas Red avidin for 2-color detection. Slides were treated with a 1:1,000 dilution of DAPI (stock 50 µg/mL) for 5 mins before mounting in Vectashield.

### Image acquisition and deconvolution

Slides from RNA and DNA FISH using fosmid derived probes were imaged on an epifluorescence microscope as previously described (Boyle et al., 2020). Step size for z stacks was set to 0.2 µm. Hardware control and image capture were performed using Nikon Nis-Elements software (Nikon) and images were deconvolved using a calculated PSF with the constrained iterative algorithm in Volocity (PerkinElmer). RNA FISH signal quantification was carried out using the quantitation module of Volocity (PerkinElmer). Number of expressing alleles were calculated by segmenting the hybridization signals and scoring each nucleus as containing 0, 1 or 2 RNA signals. DNA FISH measurements were made using the quantitation module of Volocity (PerkinElmer) and only alleles with single probe signals were analysed to eliminate the possibility of measuring sister chromatids.

Slides for DNA FISH using 10kb probes were imaged on a SoRa spinning disk confocal microscope (Nikon CSU-W1 SoRa) in 0.1 μm steps. Images were denoised and deconvolved using NIS deconvolution software (blind preset) (Nikon) and DNA FISH measurements were as for fosmid probes.

### Capture-C

The Capture-C protocol experiments was adapted from (Golov et al., 2020), with 3C libraries captured with biotin labelled fragments from BACs covering the regions of interest (Supplementary Table S7).

#### 3C *library preparation*

3x10^6^ MCF-7 cells were seeded in 10 cm plates, starved and treated as described above. Cells were crosslinked, lysed into nuclei and chromatin was fragmentated and *in situ* proximity ligated as decribed (Golov et al., 2020). For chromatin fragmentation, 350 U of DpnII 50U/µl (NEB cat. R0543M) in 300 µl 1x NEBuffer DpnII (NEB cat. B0543S) were used. After de-crosslinking and DNA extraction (Golov et al., 2020) 5 µg of DNA in 500 µl sonication buffer (25 mM Tris-HCL pH8.0, 20 mM EDTA, 0.2% SDS) were sonicated using MSE Soniprep 150 Plus at medium power for two 30 sec pulses on ice to shear DNA to a size of 150-700 bp. 1 µg of the resulting DNA was used for library preparation using the NEBNext® Ultra™ II DNA Library Prep Kit for Illumina® (NEB cat. E7103). Library PCRs were supplemented with 1× EvaGreen (Jena Bioscience cat. PCR-379). qPCRs were performed using index primers (NEBNext multiplex oligos for Illumina, set 1, E7335) and amplified to linear phase (3-5 PCR cycles).

#### Biotin-bait preparation

5 ug of pooled equimolar amounts of the selected BACs (Supplementary Table S7) were sonicated as described above for 3C libraries. Equimolarity was calculated by qPCR using primers against the chloramphenicol resistance gene (Supplementary Table S7). Forked adapters (Supplementary Table S7) were annealed in NEBuffer 2 1X (NEB cat. B7002S) to a final concentration of 20 µM in a thermal cycler set to heat at 98°C and gradually cool down to 4°C with a 1°C per 20 sec gradient.

Shredded BACs were subjected to end-preparation, A-tailing and adapter ligation as described (Golov et al., 2020). BAC baits were PCR amplified using 5’ biotinylated forward and reverse primers (Supplementary Table S7) in the following reaction mix: 10 µl 2X NEBNext® Ultra™ II Q5® Master Mix (NEB cat. M0544S), 1 µl 10 µM each bio-primer, 2 µl adapter-ligated BAC DNA, 1X EvaGreen (Jena Bioscience cat. PCR-379). The number of amplification cycles (in the linear phase of the reaction), determined in a test qPCR, usually fell between 7 and 9 cycles. Around 3 µg of biotinylated baits were obtained from 25 reactions.

#### Hybridization and biotin-pulldown

For each experiment, equal amounts of all libraries corresponding to each sample and amplified with different barcodes, were pooled and captured as a pool. One hybridization reaction was done per 1 µg of library pool. Hybridization and biotin-pulldown was done as described (Golov et al., 2020).

Singly enriched 3C libraries were amplified by qPCR using p5/p7 primers (Supplementary Table S7) as described above (Biotin-bait preparation). PCR reactions were performed with the DNA still bound to the streptavidin beads. The number of amplification cycles (in the linear phase of the reaction), determined in a test qPCR, usually fell in between 13 and 16.

A second hybridization reaction, biotin pulldown and PCR amplification was performed as before. The number of PCR cycles usually fell between 7 and 10 cycles. The pool was sequenced using either an Illumina NextSeq 500 on a single midium-output flow cell (paired-end 75-bp reads) or an Illumina Nextseq 2000 using a single P1 flow cell (paired-end 75-bp reads).

### Capture-C data analysis and generation of virtual 4C profiles

Capture-C data were aligned to the hg19 genome and processed using distiller-nf 0.3.3 (https://doi.org/10.5281/zenodo.3350937) with default settings. Coolers filtered for q>=30 were used and ICE balanced (Imakaev et al., 2012) using the cooler (Abdennur et al., 2020) command ’cooler balance --cis_only --blacklist blacklist.bed’ where blacklist.bed contains all regions outside of the capture probe area. Balanced contact frequencies were visualised using HiGlass in resgen.io (Kerpedjiev et al., 2018). Virtual 4C data were generated by extracting the balanced contact frequency values from all bins overlapping the viewpoint using cooltools (Open2C et al., 2022) and summing them. Plots were generated using seaborn (Waskom, 2021). In the case of *GREB1* locus, analysis was done at a 5 kb resolution and heatmaps and plots are shown between the coordinates chr2:11602736-11790379 (hg19). In the case of *NRIP1* locus, analysis was done at a 10 kb resolution and heatmaps and plots are shown between the coordinates chr21:16263224-16630829 (hg19).

### RT-qPCR

RNA from cells grown in 6 well plates was extracted using TRIZOL (Invitrogen 15596026) and reverse transcribed using Superscirpt II Reverse Transcriptase (Invitrogen 18064022) with random decamers. qPCR was performed using the CFX96 Touch Real-Time PCR Detection System (Bio-Rad) and the LightCycler 480 SYBR Green I Master mix (Roche 04887352001). The mean relative expression of technical replicates of each sample was measure by the CFX Manager software (Bio-Rad) according to a calibration serial dilutions curve and normalized to the mean of U2 RNA expression. Primers used are listed in Supplementary Table S5.

### Statistical analyses

DNA-FISH inter-probe distance data sets were compared using the two-tailed Mann-Whitney U test. Differences in DNA-FISH data sets comparing categorical distributions were measured using Fisher’s Exact Test. All statistical analyses were performed using R (https://www.r-project.org/).

## Data Availability

TT-seq and Capture-C data have been deposited on NCBI GEO under accession numbers GSE225508 and GSE225617 respectively.

## Author contributions

L.G.A – Conceptualization, Investigation, Methodology, Writing – original draft.

I.F. and E.T.F. – Formal analysis, Methodology.

S.B. – Investigation, Methodology.

W.A.B. – Conceptualization, Funding acquisition, Project administration, Resources, Supervision, Writing – original draft.

## Acknowledgements

We thank the Advanced Imaging Resource at the Institute of Genetics and Cancer and the Edinburgh Super-Resolution Imaging Consortium (ESRIC) for their technical support. This work has made use of the resources provided by the Edinburgh Compute and Data Facility (ECDF) (http://www.ecdf.ed.ac.uk/).

Work in the group of W.A.B. is supported by MRC University Unit grant MC_UU_00007/2. E.T.F. was supported by the Swiss National Science Foundation (P500PB_206805). Funding sources were not involved in study design, data collection, data interpretation, or the decision to submit the work for publication.

## Supplementary data

### Supplementary methods

#### Charcoal-stripped FCS

For hormone deprivation, FCS (1 litre, Gibco 10270) was charcoal-stripped of steroid hormones by heat inactivation in a waterbath at 56 °C for 30 minutes (mins) before addition of 2000U/l sulfatase (Sigma S9626-5KU, resuspended in 0.2% NaCl to 1000 U/ml). After 2 hours (hrs) incubation at 37 °C, the pH was adjusted to 4.2 using HCl and a charcoal mix (for 1 litre: 5 g charcoal (Sigma-Aldrich 05105), 25 mg dextran T70 (USP, 1179741), 50 ml water) was added and incubated overnight (o/n) at 4 °C with stirring. Charcoal was removed by centrifugation at 500 *g* for 30 mins at 4 °C, the pH re-adjusted to 4.2 and a second charcoal mix added, incubated and then removed. Centrifugation was repeated to remove residual charcoal and the pH adjusted to 7.2 with NaOH. Stripped FCS was filter sterilized, aliquoted and stored at -20°C.

#### CRISPR deletions

pSpCas9(BB)-2A-GFP (PX458, Addgene 48138) plasmids including either the left or right gRNA (Supplementary Table S5) were co-transfected into MCF-7 cells using Lipofectamine 3000 (Invitrogen, L3000015). After 24 hrs, single GFP+ve cells were seeded into 96 well plates. Cell lysates from different clones were obtained using DirectPCR Lysis Reagent (Viagen Biotech, 102-T) and genotyping was performed by PCR using the primers in Supplementary Table S5. PCR bands were cloned into the TOPO-TA cloning kit (Invitrogen, 450071) and validated by Sanger sequencing.

#### Immunofluorescence

4x10^5^ MCF-7 cells were seeded on Thermo Scientific SuperFrost Plus Adhesion slides (15438060), starved and treated as specified above. Before fixing, cells were rinsed with 0.5% Triton X-100/ PBS for a few seconds and then slides were fixed with pFa 4% in PBS for 10 min, rinsed with PBS 3 times for 2 min and permeabilized with 0.5% Triton X-100/ PBS for 10 min. Following 3x 2 min PBS rinses, slides were blocked with 1% BSA in PBS for 30 min at r.t. and incubated with 1:250 mouse anti ERα antibody (F-10, Santa Cruz, cat. Sc-8002) in 1% BSA in PBS for 1h. Slides were rinsed twice in PBS for 2 min and incubated for 45 min with 1:500 Goat anti-Mouse Alexa Fluor 488 antibody (Invitrogen, cat. A11001) in 1% BSA/ PBS. After rinsing, slides were incubated with a 1:1,000 dilution of DAPI (stock 50 µg/mL) for 5 mins before mounting in Vectashield (Vector Laboratories, cat. H1000).

Slides for immunofluorescence were imaged on an epifluorescence microscope (Zeiss AxioImager 2, camera Hamamatsu Orca Flash 4.0 16-bit) and nuclear fluorescence intensity was quantified using CellProfiler v3.1.8.

**Supplementary Figure S1.**
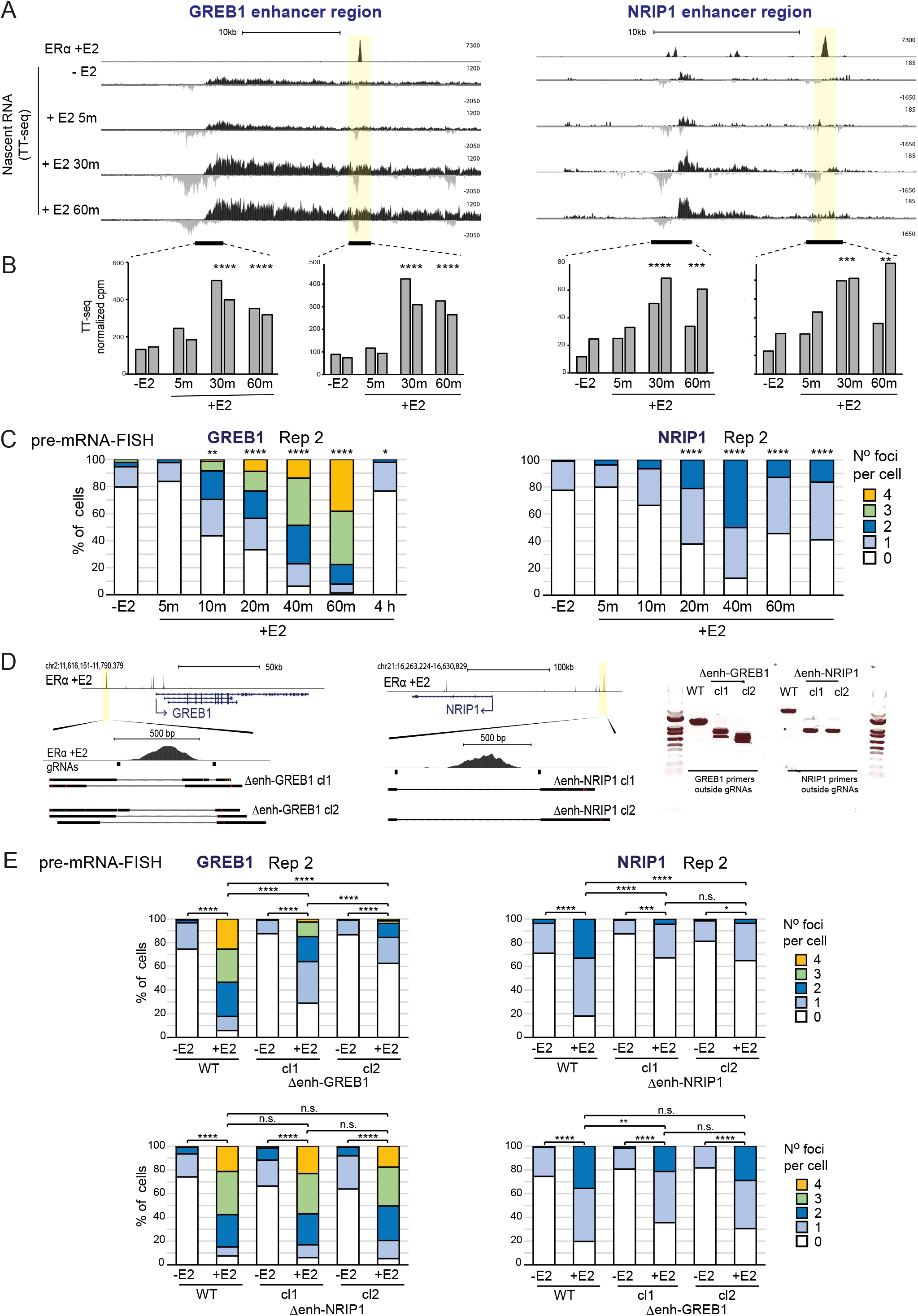
Related to Figure 1. Rapid induction of transcription at ER-responsive enhancers in MCF-7 cells. A) Genome Browser screen shots showing ChlP-seq tracks of ERa after E2 addition and TT-seq tracks without E2 and 5, 30 and 60 min after E2 addition over the intergenic regions around the *GREB1* and *NRIP1* putative enhancers. B) Quantification of TT-seq reads over the indicated regions. These correspond to ATAC-seq peak regions extended for 1,5 kb at either side. Normalized counts per million reads (cpm) of two replicates are shown. **p<0.01, ***p<0.001, ****p<0.0001. C) Biological replicate of the data in Fig. 1C. Pre-mRNA FISH for *GREB1* and *NRIP1* nascent transcripts without and with E2 in MCF-7 cells for the indicated timepoints. The percentage of cells with 0, 1, 2, 3, 4 foci at each timepoint is shown. Two­sided Fisher exact test, *p<0.05, **p<0.01, ***p<0.001, ****p<0.0001. D) Genome browser screens shots of (left) *GREB1* and (centre) *NRIP1* loci including ERa +E2 ChIP seqtrack (Zwart et al, 2011) indicating the position of gRNAs used in CRISPR-Cas9 enhancer deletions. Aligned sequences of the homozygous clones for each deletion are shown below. Right; agarose gel of PCRs products from genomic DNA extracted from the different deletion clones using primers around the deletions. E) Upper panels; biological replicate of data in Fig. 1D. pre-mRNA FISH for *GREB1* and *NRIP1* nascent transcripts in WT MCF-7 cells and in cells where the respective putative ERa enhancers have been deleted. The percentage of cells with 0, 1, 2, 3 or 4 foci of each cell line, without and with E2 is shown. Results for two clones bearing each of the deletions in homozygosity are shown. Lower panels; as in upper panels but assaying for *GREB1* nascent transcripts in in cells deleted for the NRIP1 enhancers (left) and for (right) *NRIP1* nascent transcripts in cells deleted for the GREB1 enhancer. Two-sided Fisher exact test. n.s. not significant, *p<0.05, **p<0.01, ***p<0.001, ****p<0.0001. Statistical data are in Supplementary Table S1

**Supplementary Figure S2.**
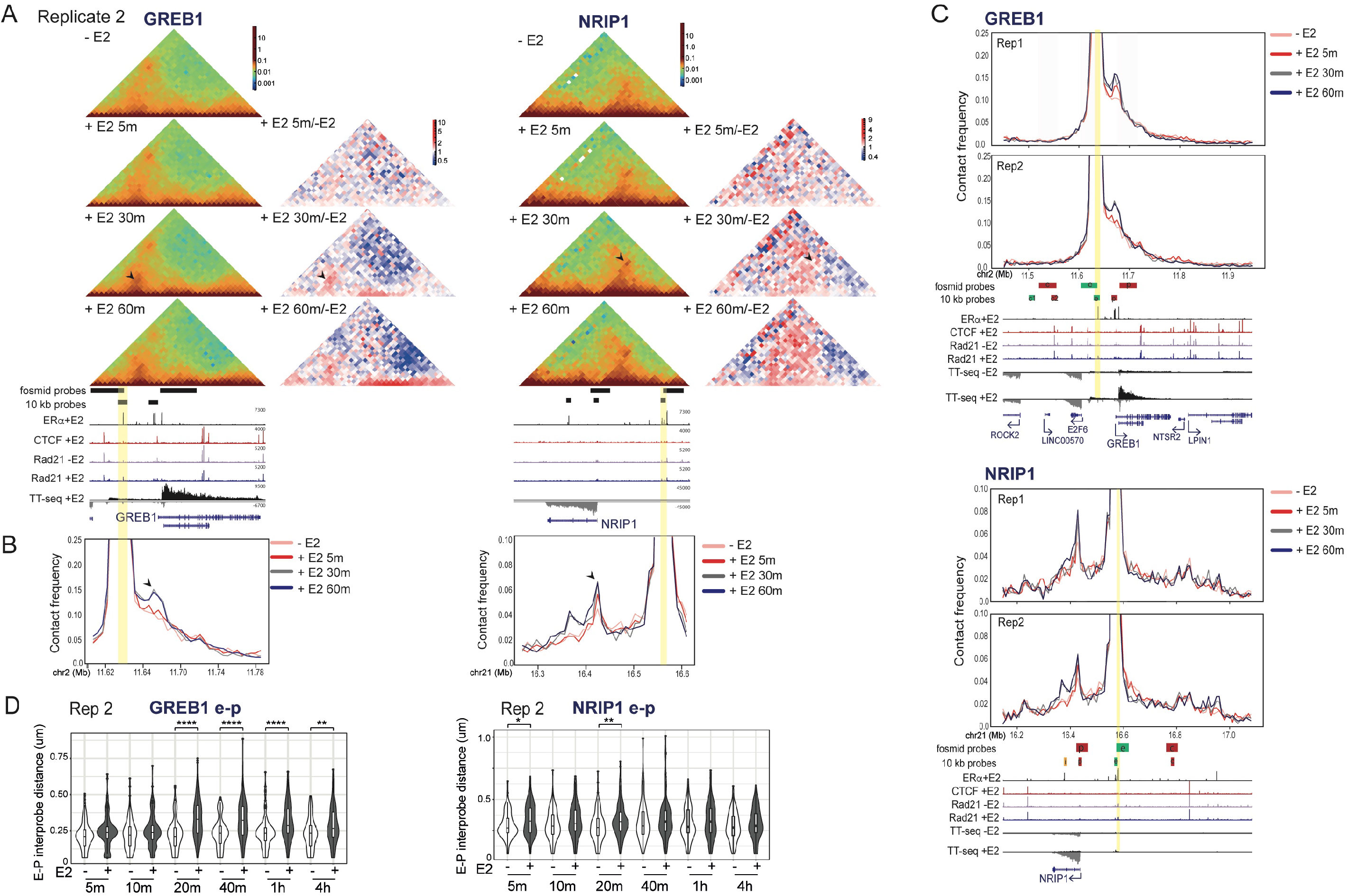
Related to Figure 2. A and B) Biological replicates for the data in Fig. 2A,B. C) Virtual 4C plots derived from the Capture-C normalized contact frequencies for the whole of the captured regions using the enhancer regions marked with a yellow bar as viewpoints. The relative positions and size of the probes used in DNA-FISH are shown below along with ERa +E2 ChlP-seq (Zwart et al, 2011), CTCF +E2, Rad21 -E2, Rad21 +E2 ChlP-seq (Schmidt et al, 2010), and TT-seq +E2 30 min are also shown. Data shown for both biological replicates. Genome coordinates: hg19 human genome. D) Biological replicates for the data in Fig. 2D. The statistical difference in data distribution +/-E2 at each time point was assessed by a two-sided Mann-Whitney test. *p<0.05, **p<0.01, “*p<0.001, “**p<0.0001. Holm-Bonferroni correction for multiple testing. Statistical data are in Supplementary Table S2.

**Supplementary Figure S3.**
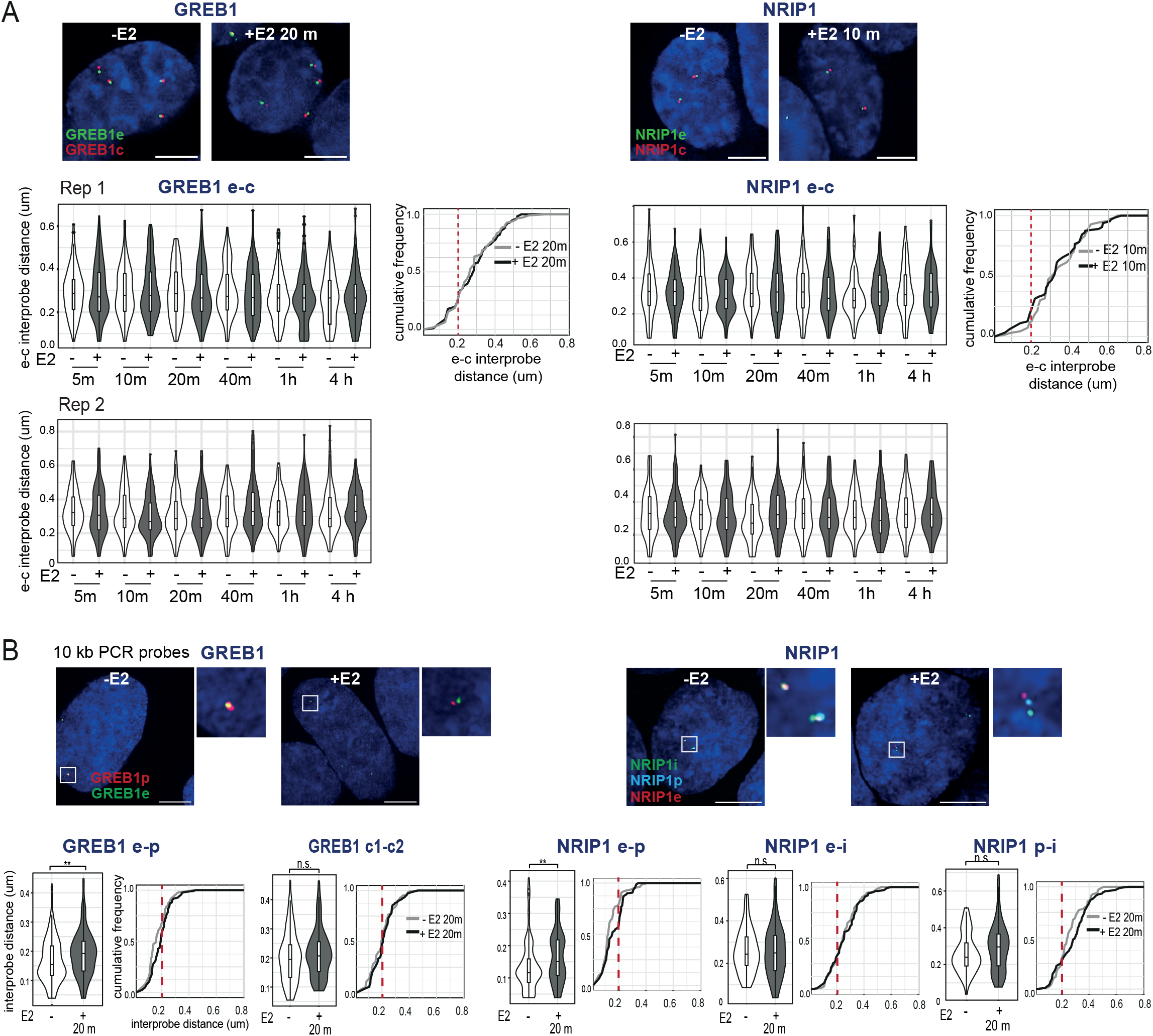
Related to Figure 2. A) Top; Representative images of nuclei (DAPI, blue) of MCF-7 cells untreated (-E2) or treated with E2 (+E2) for the indicated time points showing DNA-FISH signal from fosmid probes targeted to the enhancer (e, green) and control (c, red) regions of either GREB1 or NRIP1 loci. Scale bars, 5pm. Below; Violin and cumulative frequency plots showing the distribution of DNA-FISH interprobe distances in untreated (-E2) and E2 (+E2) treated MCF-7 cells for the indicated time points and using e-c fosmid probe pairs. Boxplots indicating the median distances are included. In cumulative frequency plots the red dashed lines indicate 200 nm distance. Data shown for two biological replicates. B) Top; Representative images of nuclei (DAPI, blue) of MCF-7 cells untreated (-E2) or treated with E2 (+E2) for 30 min showing DNA-FISH signal from 10 kb probes targeted to the *GREB1* enhancer (GREBIe), promoter (GREBIp), control (GREB1c1 and GREB1c2), and the *NRIP1* promoter (NRIPIp), enhancer (NRIPIe) and intragenic (NRIP1I) regions. Scale bars, 5pm. Below; Violin and cumulative frequency plots showing the distribution of DNA-FISH interprobe distances between GREB1 e-p, GREB1 c1-c2, NRIP1 e-p, NRIP1 e-i, NRIP1 p-i 10 kb probe pairs in MCF-7 cells with and without E2. Statistical significance of data distributions were assayed using a Two-sided Mann-Whitney test. n.s. p> 0.05, **p<0.01. Statistical data are in Supplementary Table S2.

**Supplementary Figure S4.**
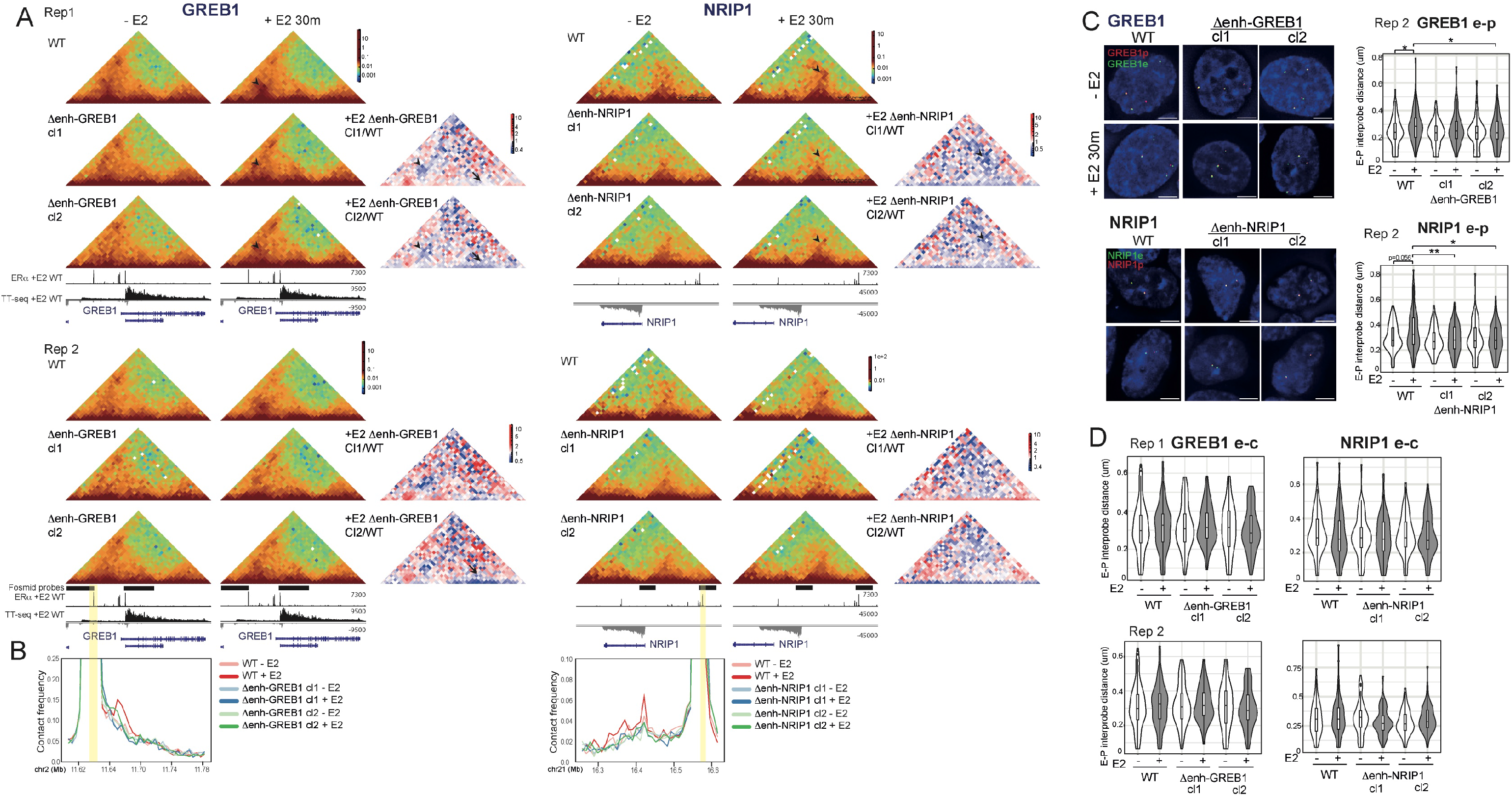
Related to Figure 2. A) Capture-C heatmaps at *GREB1* (left; 5kb resolution) and *NRIP1* (right; 10kb resolution) loci from WT and *GREB1* or *NRIP1* enhancer deletion clones without (-E2) and with E2 (+E2) for 30 min. ERa +E2 (Zwart et al, 2011) and TT-seq +E2 30 min and Gene tracks are shown below. In red-green-blue heatmaps, each pixel represents the normalized contact frequency between a pair of loci. In red-white-blue heatmaps each pixel represents the ratio between E2-treated enhancer deletion clones vs E2-tretead WT samples. Blue pixels indicate loss of contact frequency in enhancer deletion vs WT samples and red pixels indicate a gain. Black arrowheads indicate pixels corresponding to the enhancer-promoter pairs whose E2-dependent contact frequency is lost in enhancer deletion cells. Arrows indicate the loss of very short contact frequencies due to the reduction in transcription at *GREB1* in enhancer deletion clones in contrast to WT cells. Genome coordinates: hg19 human genome. BVirtual 4C plots derived from the Capture-C normalized contact frequencies for replicate 2 from panel (A) using the enhancer regions marked with a yellow bar as viewpoints. C) Left; Representative images of nuclei (DAPI, blue) of WT or enhancer deletion MCF-7 clones treated as indicated showing DNA-FISH signal from fosmid probes targeted to the enhancer (e, green), and the promoter (p, red)or (right) control (c, red) regions of the GREB1 or NRIP1 loci. Scale bars, 5pm. Right; Violin plots showing the distribution of DNA-FISH interprobe distances between e-p probes in WT or enhancer deletion MCF-7 clones untreated or treated with E2 for 30 min. Boxplots indicating the median distances are included. Two-sided, Mann-Whitney test, Holm-Bonferroni correction for multiple testing. Biological replicate for the data in Figure 2F. D) Violin plots as for (C) but with e-c fosmid probe pairs for two replicate experiments. *p<0.05, **p<0.01. Statistical data are in Supplementary Table S2.

**Supplementary Figure S5.**
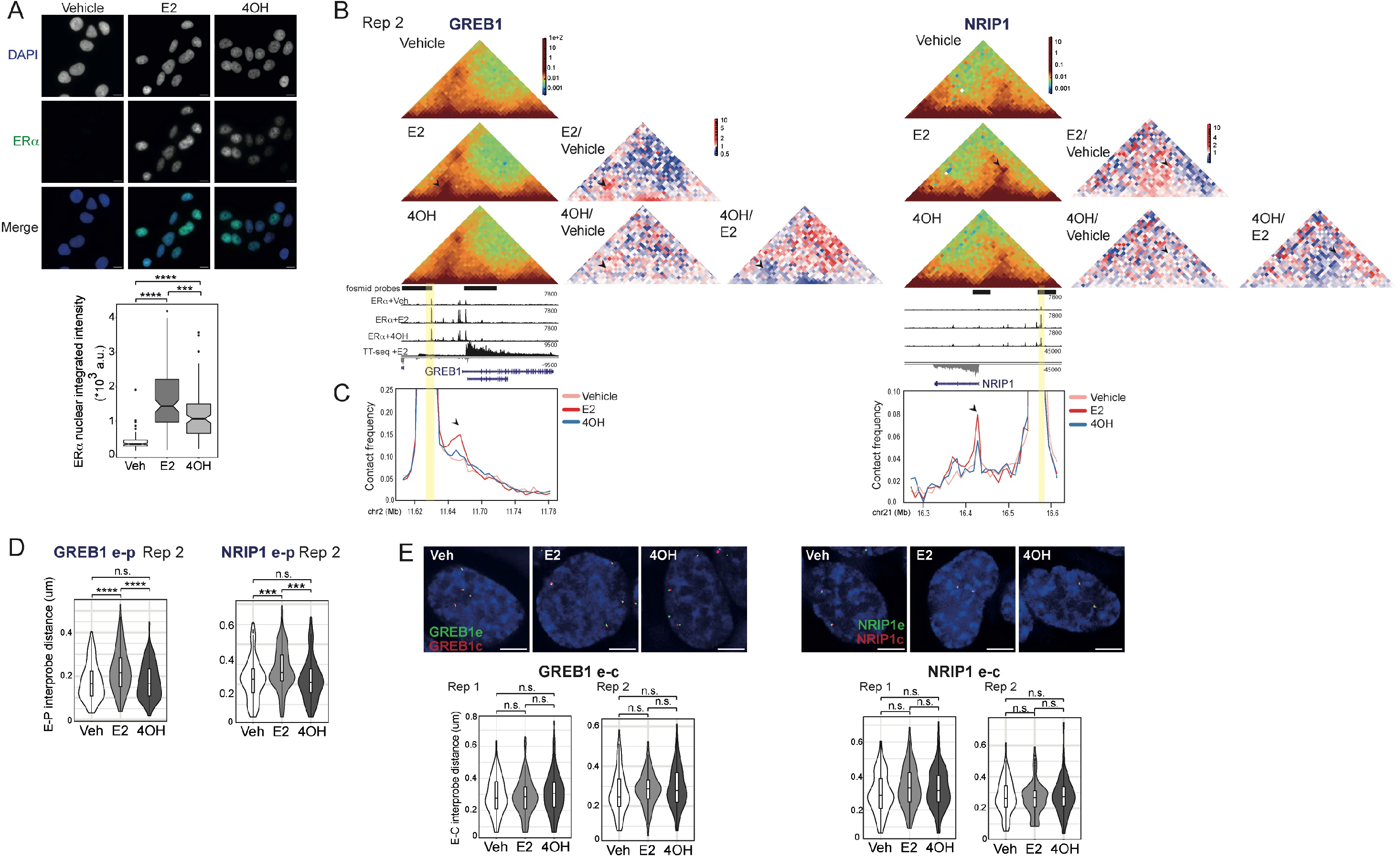
Related to Figure 3. A) Representative images of nuclei (DAPI, blue in merge) subject to immunofluorescence for ERa (green in merge) in MCF-7 cells treated with vehicle, E2 or 4OH for 30 min and pre-extracted with 0.5% Triton X-100 before fixation. Scale bar, 10 pM. Below, boxplots showing the integrated nuclear intensity. Two-sided Mann-Whitney test, Holm-Bonferroni correction for multiple testing. B and C) Biological replicate of Capture-C data in Fig. 3B,C. D) Violin plots showing the distribution of DNA-FISH interprobe distances of cells treated with vehicle, E2 or 4OH using e-p fosmid probes at either *GREB1* or *NRIP1* loci. Boxes indicate the median and interquartile distances. Two-sided Mann-Whitney test, Holm-Bonferroni correction for multiple testing, n.s. p>0.05, *“p<0.001, ****p<0.0001. Biological replicate for the data in Fig. 3E. E) Top;DNA-FISH of MCF-7 cells treated with vehicle, E2 or 4OH for 30 min using fosmid probes targeted to the enhancer (e) and control (c) regions of either *GREB1* or *NRIP1* loci. Scale bars, 5pm. Below; As in (D) but for enhancer (e) and control (c) probe pairs. Two biological replicates. Statistical data are in Supplementary Table S3.

**Supplementary Figure S6.**
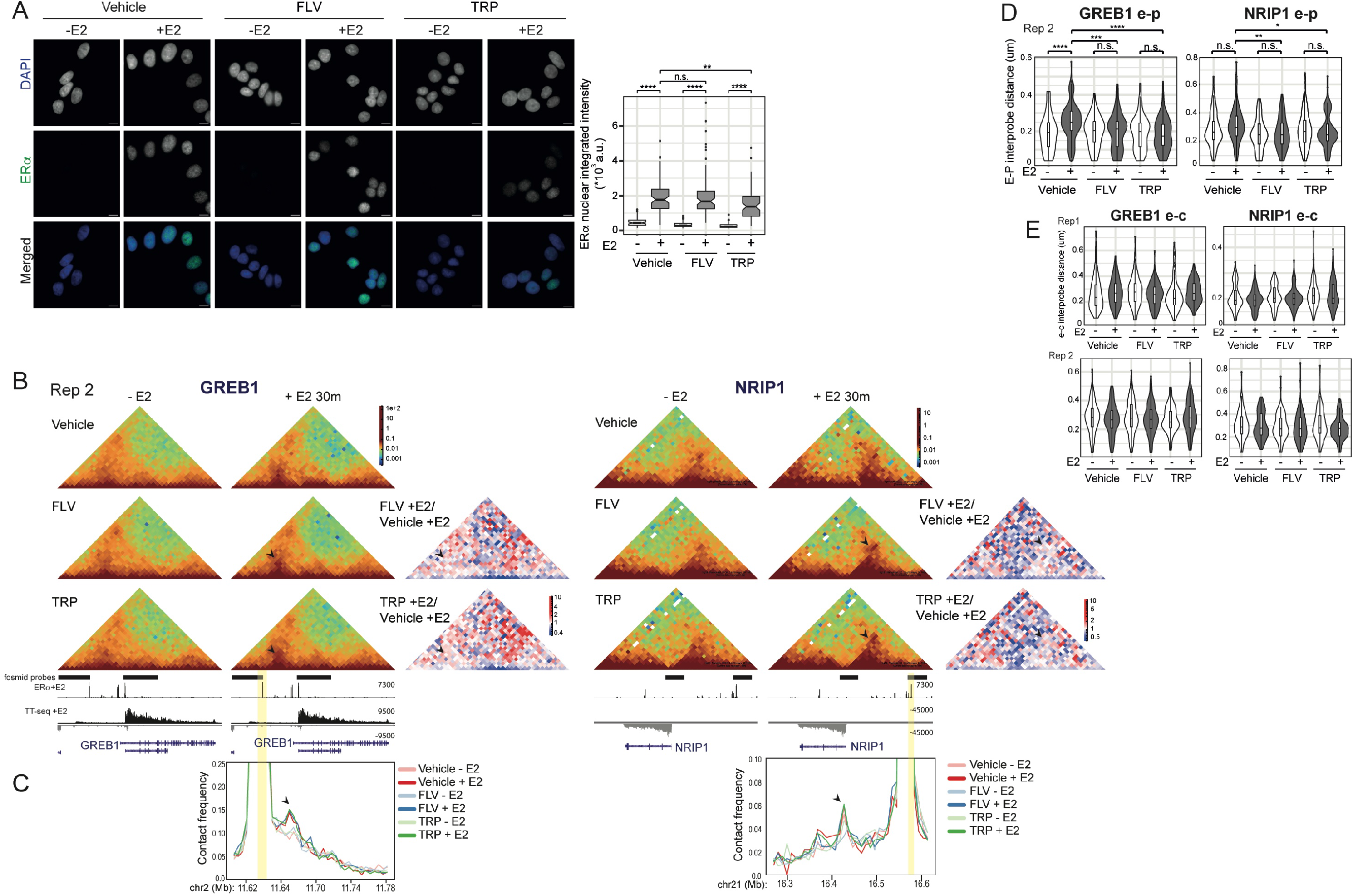
Related to Figure 4. **A)** Left: representative images of nuclei (DAPI, blue in merged) subject to immunofluorescence for ERa (green in merge) in MCF-7 cells treated with vehicle, E2 or4OH for 30 min and pre-extracted with 0.5% Triton-X-100 before fixing. Scale bar, 10 pM. Right: boxplots showing the integrated nuclear intensity. Two-sided Mann-Whitney test, Holm-Bonferroni correction for multiple testing. B and C) Biological replicate of data in Fig. 4B and C. D) Violin plots showing the distribution of DNA-FISH enhancer-promoter interprobe distances at the *GREB1* or *NRIP1* loci in cells treated as indicated. Boxes indicating the median distances. Two-sided Mann-Whitney test, Holm-Bonferroni correction for multiple testing. Biological replicate for the data in Fig. 4E. n.s p> 0.05, *p<0.05, **p<0.01, ***p<0.001, ****p<0.0001. E) Violin plots showing the distribution of DNA-FISH enhancer-control interprobe at *GREB1* or *NRIP1* loci for cells treated with vehicle, FLV or TRP for 5 min prior to treatment without (-E2) and with E2 (+E2) for 30 min. Boxes indicating the median distances. Two-sided Mann-Whitney test, Holm-Bonferroni correction for multiple testing shows no significant differences. Data from two biological replicates shown. Statistical data for are in Supplementary Table S4.

## References

Abdennur N, Mirny L. A. 2020.Cooler: Scalable storage for Hi-C data and other genomically labeled arrays. Bioinformatics 36: 311–316.

Alexander JM, Guan J, Li B, Maliskova L, Song M, Shen Y, Huang B, Lomvardas S, Weiner OD. 2019.Live-cell imaging reveals enhancer-dependent *Sox2* transcription in the absence of enhancer proximity. Elife 8: e41769.

Barshad G, Lewis JJ, Chivu AG, Abuhashem A, Krietenstein N, Rice EJ, Ma Z, Rando OJ, Hadjantonakis AK, Danko CG. 2023. RNA polymerase II dynamics shape enhancer-promoter interactions. Nat Genet. doi:10.1038/s41588-023-01442-7.

Benabdallah NS, Williamson I, Illingworth RS, Kane L, Boyle S, Sengupta D, Grimes GR, Therizols P, Bickmore WA. 2019. Decreased enhancer-promoter proximity accompanying enhancer activation. Mol. Cell 76: 473–484.e7.

Bensaude O. 2011. Inhibiting eukaryotic transcription: Which compound to choose? How to evaluate its activity? Transcription 2: 103–108.

Boyle S, Flyamer IM, Williamson I, Sengupta D, Bickmore WA, Illingworth RS. 2020. A central role for canonical PRC1 in shaping the 3D nuclear landscape. Genes Dev. 34: 931–949.

Chen F, Gao X, Shilatifard A, Shilatifard A. 2015. Stably paused genes revealed through inhibition of transcription initiation by the TFIIH inhibitor triptolide. Genes Dev. 29: 39–47.

Chen H, Levo M, Barinov L, Fujioka M, Jaynes JB, Gregor T. 2018. Dynamic interplay between enhancer-promoter topology and gene activity. Nat Genet. 50:1296–1303.

Finn EH, Pegoraro G, Brandão HB, Valton AL, Oomen ME, Dekker J, Mirny L, Misteli T. 2019. Extensive heterogeneity and intrinsic variation in spatial genome organization. Cell 176: 1502–1515.e10.

Fudenberg G, Imakaev M. 2017. FISH-ing for captured contacts: towards reconciling FISH and 3C. Nat Methods 14: 673–678.

Fullwood MJ, Liu MH, Pan YF, Liu J, Xu H, Mohamed YB, Orlov YL, Velkov S, Ho A, Mei PH, et al. 2009. An oestrogen-receptor-alpha-bound human chromatin interactome. Nature. 462: 58–64.

Gavrilov A, Razin SV, Cavalli G. 2015. In vivo formaldehyde cross-linking: It is time for black box analysis. Briefings in Functional Genomics 1:, 163–165.

Giorgetti L, Heard E. 2016. Closing the loop: 3C versus DNA FISH. Genome Biology 17: 215.

Glont SE, Chernukhin I, Carroll JS. 2019. Comprehensive genomic analysis reveals that the pioneering function of FOXA1 Is Independent of hormonal signaling. Cell Reports 26: 2558–2565.e3.

Golov AK, Ulianov SV, Luzhin AV, Kalabusheva EP, Kantidze OL, Flyamer IM, Razin SV, Gavrilov AA. 2019. C-TALE, a new cost-effective method for targeted enrichment of Hi-C/3C-seq libraries. Methods 170: 48–60.

Guan J, W. Zhou M, Hafner RA, Blake C, Chalouni IP, Chen, De Bruyn JM, Giltnane SJ, Hartman A, Heidersbach R. 2019.Therapeutic Ligands Antagonize Estrogen Receptor Function by Impairing Its Mobility. Cell 178: 949–963.

Hah N, Danko CG, Core L, Waterfall JJ, Siepel A, Lis JT, Kraus WL. 2011. A rapid, extensive, and transient transcriptional response to estrogen signaling in breast cancer cells. Cell 145: 622–634.

Hah N, Murakami S, Nagari A, Danko CG, Kraus WL. 2013. Enhancer transcripts mark active estrogen receptor binding sites. Genome Res. 23: 1210–1223.

Hoffman EA, Frey BL, Smith LM, Auble DT. 2015. Formaldehyde crosslinking: A tool for the study of chromatin complexes. J. Biol. Chem. 290: 26404–26411.

Holding AN, Cullen AE, Markowetz F. 2018. Genome-wide Estrogen Receptor-α activation is sustained, not cyclical. eLife. 7: e40854.

Hsieh TS, Cattoglio C, Slobodyanyuk E, Hansen AS, Rando OJ, Tjian R, Darzacq X. 2020. Resolving the 3D landscape of transcription-linked mammalian chromatin folding. Mol Cell. 78: 539–553.e8.

Hurtado A, Holmes KA, Ross-Innes CS, Schmidt D, Carroll JS. 2011.FOXA1 is a key determinant of estrogen receptor function and endocrine response. Nat. Genet. 43: 27–33.

Imakaev M, Fudenberg G, McCord RP, Naumova N, Goloborodko A, Lajoie BR, Dekker J, Mirny LA. 2012. Iterative correction of Hi-C data reveals hallmarks of chromosome organization. Nat. Methods 9: 999–1003.

Irgen-Gioro S, Yoshida S, Walling V, Chong S. 2022.Fixation can change the appearance of phase separation in living cells. Elife 11: e79903.

Javierre BM, Sewitz S, Cairns J, Wingett SW, Várnai C, Thiecke MJ, Freire-Pritchett P, Spivakov M, Fraser P, Burren OS, et al. 2016. Lineage-Specific Genome Architecture Links Enhancers and Non-coding Disease Variants to Target Gene Promoters. Cell 167: 1369–1384.

Jonkers I, Kwak H, Lis JT. 2014. Genome-wide dynamics of Pol II elongation and its interplay with promoter proximal pausing, chromatin, and exons. eLife 3: e02407.

Jubb AW, Boyle S, Hume DA, Bickmore WA. 2017. Glucocorticoid receptor binding induces rapid and prolonged large-scale chromatin decompaction at multiple target loci. Cell Rep. 21: 3022–3031.

Kane L, Williamson I, Flyamer IM, Kumar Y, Hill RE, Lettice LA, Bickmore WA. 2022. Cohesin is required for long-range enhancer action at the Shh locus. Nat Struct Mol Biol. 29: 891–897.

Karr JP, Ferrie JJ, Tjian R, Darzacq X. 2022. The transcription factor activity gradient (TAG) model: contemplating a contact-independent mechanism for enhancer-promoter communication. Genes Dev. 36: 7–16.

Kerpedjiev P, Abdennur N, Lekschas F, McCallum C, Dinkla K, Strobelt H, Luber JM, Ouellette SB, Azhir A, Kumar N, et al. 2018. HiGlass: Web-based visual exploration and analysis of genome interaction maps. Genome Biology 19: 125.

Kocanova S, Kerr EA, Rafique S, Boyle S, Katz E, Caze-Subra S, Bickmore WA, Bystricky K. 2010. Activation of estrogen-responsive genes does not require their nuclear co-localization. PLoS Genet. 6: e1000922.

Lee JH, Wang R, Xiong F, Krakowiak J, Liao Z, Nguyen PT, Moroz-Omori EV, Shao J, Zhu X, Bolt MJ, et al. 2021.Enhancer RNA m6A methylation facilitates transcriptional condensate formation and gene activation. Mol Cell 81: 3368–3385.e9.

Légaré S, Basik M. 2016.The Link Between ERα Corepressors and Histone Deacetylases in Tamoxifen Resistance in Breast Cancer. Mol Endocrinol. 30: 965–976.

Leidescher S, Ribisel J, Ullrich S, Feodorova Y, Hildebrand E, Galitsyna A, Bultmann S, Link S, Thanisch K, Mulholland C, *et a*l. 2022.Spatial organization of transcribed eukaryotic genes. Nat. Cell Biol. 24: 327–339.

Leung AK, Yao L, Yu H. 2022.Functional genomic assays to annotate enhancer-promoter interactions genome wide. Hum Mol Genet. 31: R97–R104.

Li W, Notani D, Ma Q, Tanasa B, Nunez E, Chen AY, Merkurjev D, Zhang J, Ohgi K, Song X, et al. 2013.Functional roles of enhancer RNAs for oestrogen-dependent transcriptional activation. Nature 498: 516–520.

Lim B, Levine MS. 2021. Enhancer-promoter communication: hubs or loops? Curr Opin Genet Dev. 67: 5–9.

McCord RP, Kaplan N, Giorgetti L. 2020.Chromosome conformation capture and beyond: Toward an integrative view of chromosome structure and function. Mol Cell 77: 688–708.

Mateo LJ, Murphy SE, Hafner A, Cinquini IS, Walker CA, Boettiger AN. 2019. Visualizing DNA folding and RNA in embryos at single-cell resolution. Nature. 568: 49–54.

Open2C, Abdennur N, Abraham S, Fudenberg G, Flyamer IM, Galitsyna AA, Goloborodko A, Imakaev M, Oksuz BA, Venev SV. 2022. Cooltools : enabling high-resolution Hi-C analysis in Python. (iv). bioRxiv doi: https://doi.org/10.1101/2022.10.31.514564.

Platania A, Erb C, Barbieri M, Molcrette B, Grandgirard E, de Kort MA, Meaburn K, Taylor T, Shchuka VM, Kocanova S, et al. (2023) Competition between transcription and loop extrusion modulates promoter and enhancer dynamics. bioRxiv 2023.04.25.538222.

Rinzema NJ, Sofiadis K, Tjalsma SJD, Verstegen MJAM, Oz Y, Valdes-Quezada C, Felder AK, Filipovska T, van der Elst S, de Andrade Dos Ramos Z, et al. (2022). Building regulatory landscapes reveals that an enhancer can recruit cohesin to create contact domains, engage CTCF sites and activate distant genes. Nat Struct Mol Biol. 29: 563–574.

Rodriguez J, Ren G, Day CR, Zhao K, Chow CC, Larson DR. 2019. Intrinsic dynamics of a human gene reveal the basis of expression heterogeneity. Cell 176: 213–226.e18.

Saravanan B, Soota D, Islam Z, Majumdar S, Mann R, Meel S, Farooq U, Walavalkar K, Gayen S, Singh AK, et al. 2020. Ligand dependent gene regulation by transient ERα clustered enhancers. PLoS Genet. 16: e1008516.

Schwalb B, Miche, M, Zacher B, Frühauf K, Demel C, Tresch A, Gagneur J, Cramer P. 2016. TT-seq maps the human transient transcriptome. Science 352: 1225–1228.

Schmidt D, Schwalie PC, Ross-Innes CS, Hurtado A, Brown GD, Carroll JS, Flicek P, Odom DT. 2010. A CTCF-independent role for cohesin in tissue-specific transcription. Genome Res. 20: 578–588.

Schmiedeberg L, Skene P, Deaton A, Bird A. 2009. A temporal threshold for formaldehyde crosslinking and fixation. PLoS One. 4, e4636.

Teves SS, An L, Hansen AS, Xie L, Darzacq X, Tjian R. 2016. A dynamic mode of mitotic bookmarking by transcription factors. Elife. 5: e22280.

Waskom ML. 2021. seaborn: statistical data visualization. J Open Source Software 6: 3021.

Williamson I, Berlivet S, Eskeland R, Boyle S, Illingworth RS, Paquette D, Dostie J, Bickmore WA. 2014. Spatial genome organization: contrasting views from chromosome conformation capture and fluorescence in situ hybridization. Genes Dev. 28: 2778–2791.

Zwart W, Theodorou V, Kok M, Canisius S, Linn S, Carroll JS. 2011. Oestrogen receptor-co-factor-chromatin specificity in the transcriptional regulation of breast cancer. EMBO J. 30: 4764–4776.

